# Should I stay or should I go? Modelling the decision-making process behind ungulate partial migration

**DOI:** 10.64898/2026.07.07.737075

**Authors:** Joel O. Abraham, Ricardo Martinez-Garcia, Finote Gijsman, Erin M. Phillips, Corina E. Tarnita

## Abstract

Despite the ecological importance of ungulate migrations, we lack a complete understanding of why some ungulates migrate and others do not. Though progress has been made towards understanding differences across species and between populations, migratory behavior varies even *within* populations: in many populations, some individuals remain behind as residents (partial migration). Theoretical population-level work has suggested that these different migratory tactics can coexist, but such approaches stop short of providing insights into how individuals make the decision to stay or go each year. Using long-term data from three ungulate populations, we find that individuals’ probabilities of migrating are highly variable across years, which points to a non-trivial context-dependent decision-making process, whose underlying mechanisms must be probed via individual-level modeling. Drawing on existing knowledge, we propose a decision-making model of ungulate migration onset wherein individuals probabilistically decide to start migrating based on the local intensity of environmental and/or social cues. Residents arise as a robust collective organization phenomenon in our model. At sufficiently large population sizes, the number of residents is invariant with total population size, consistent with empirical patterns. Instead, resident numbers are influenced by the severity of the ‘bad’ season, by relevant character differences among individuals, and by how individuals contribute and respond to environmental and/or social cues; for instance, when social cues contribute to decision-making in addition to environmental ones, fewer residents result, and migration is more likely to be complete. Overall, our model provides a potential mechanistic explanation for how residents might emerge within migratory ungulate populations.

## INTRODUCTION

Ungulate migration is a socially and ecologically important phenomenon. Mass migrations of ungulates, such as that of the Serengeti wildebeest (*Connochaetes taurinus*), have long captivated public imagination (1, 2), and consequently generate revenue for local economies via international attention and ecotourism (3). Migration helps ungulates cope with heterogeneous environments (4, 5), supports large populations (6–8), and couples ecosystem processes across vast spatial scales (9). However, migratory behavior varies hugely within ungulates, and we still do not fully understand why some ungulates migrate and others do not (10–12).

Significant progress has been made towards explaining differences in migratory behavior across species and between populations. At the species and population scales, variation in migration has been linked to environmental, social, and species-specific factors, such as differences in rates of vegetation green-up (13, 14), accumulated cultural knowledge (15, 16), and species characteristics like body size and grass dependence (14, 17). However, migration varies even within a population, where species characteristics, social context, and environmental features are held constant: in many populations of migratory animals, a fraction of the population does not partake in the migration, instead remaining behind as ‘residents’ (12, 18–21).

The mechanisms underlying the existence of residents remain a persistent unknown of migration, within ungulates and beyond (10–12, 20, 21). One proposed hypothesis for this phenomenon is that the factors that explain differences in migration at other scales—*e.g.*, differences in body size—might also explain within-population variation in migration. Consequently, differences in various characteristics of migrants and residents within partially migratory populations have been widely documented (12, 22–27), and yet character differences do not always mirror species-level patterns. For example, while migratory behavior is more common among large-bodied species (5), migrants tend to be smaller on average than residents within the Serengeti wildebeest population (26). Moreover, differences between migrants and residents vary across populations: in some populations, older individuals are more likely to migrate, while younger individuals are more migratory in others (12). As such, character differences between migrants and residents are not sufficient to explain the existence of residents.

A related hypothesis is that the individuals that stay behind are genetically distinct from those that migrate, perhaps lacking some genetic architecture that would allow for migration (10, 12, 20, 21, 28). Although such a hypothesis is, in general, hard to evaluate exhaustively, genetic surveys on a diversity of ungulate populations have failed to find consistent genetic differences between migrant and resident individuals (12, 29–34). Even more compellingly, robust long-term monitoring efforts and the expansion of GPS tracking technologies have revealed that individuals readily switch between migratory tactics, migrating in one year and remaining as residents in the next. A recent synthesis found evidence of migratory switching in 20 out of the 23 ungulate populations for which data were available (12), and migratory switching has been recorded even within the Serengeti wildebeest (35). In other words, it is unlikely that migrants and residents are entirely genetically distinct subpopulations.

Building on this updated understanding, recent work has employed multi-year population-level modeling where migration is treated as a plastic phenotype and individuals are assumed to choose migratory tactics (*i.e.,* resident versus migrant) probabilistically (18, 19, 36). This work has demonstrated that there are indeed conditions under which the different tactics yield equal fitness for individuals, allowing the two tactics to stably coexist in accordance with the Ideal Free Distribution. However, because these models focus on the population-level outcome rather than the individual decision-making process, the mechanism that underlies the choice of migratory tactic remains an important open question. The simplest imaginable mechanism would be a genetically fixed stochastic switch, honed by evolution over long timescales. Under such a mechanism, migration would operate like a population-level ‘coin toss’, wherein every individual has the same genetically determined probability of migrating. Because this probability is fixed, the fraction of the population that migrates would remain roughly constant from year to year, changing only gradually across generations. Consequently, both migrant and resident numbers would increase linearly with total population size, with larger total populations simply producing more individuals of both types. An alternative, albeit more complex, decision-making mechanism would be a context-dependent switch: *i.e.*, an individual’s probability of migrating would not be genetically fixed, but instead would adjust dynamically in response to environmental, and potentially also social, cues. If this is the case, then observed migration probabilities could vary yearly as environmental and social contexts change. Consequently, resident and migrant numbers might not scale linearly with total population size, but rather exhibit complex, potentially nonlinear relationships. Unlike in the previous case of a genetically fixed migration probability, understanding this context-dependent decision-making requires a mechanistic model to explore how individuals integrate and weigh different sources of information when deciding whether to migrate.

Here, we compile abundance data from three different ungulate populations—Banff elk (*Cervus canadensis*), Mongolian gazelle (*Procapra gutturosa*), and Serengeti wildebeest (*Connochaetes taurinus*)—and evaluate whether individuals migrate with some probability *p* that is constant or variable over the multidecadal timescales of these datasets. We find pronounced year-to-year variation in migration probabilities that is inconsistent with a genetically fixed migration probability (**Fig. 1**). Consistently, we find that the number of migrants does scale linearly with the total population size, whereas total population size cannot explain variation in the number of residents (**Fig. 1**), implying that the decision to migrate is context-dependent. Consequently, the decision-making process of individuals must be modeled explicitly to understand how the population-level partitioning into migrants and residents occurs each year. To this end, we propose a bottom-up decision-making model of migration onset that incorporates empirical insights into how migratory ungulates decide to migrate based on environmental and social cues (**Fig. 2**). With this model, we investigate the conditions that lead to the existence of a resident subpopulation and the potential roles of both environmental and social information in driving and/or amplifying migration; simultaneously, we explore the extent to which such a context-dependent decision-making process can account for empirical patterns of partial migration.

**FIGURE 1.**
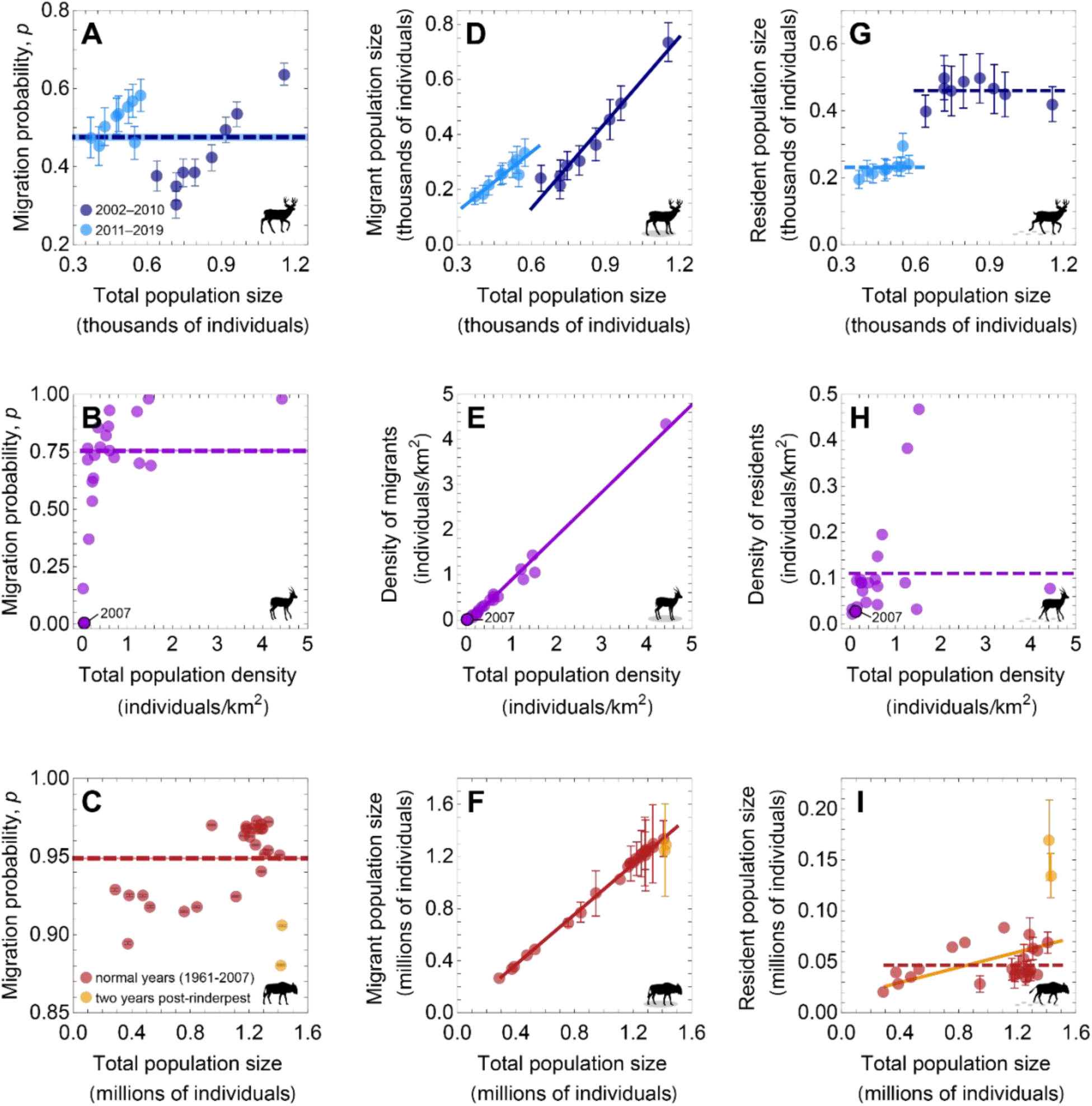
Empirical patterns of ungulate partial migration. Populations include: (**A, D, G**) elk (*Cervus canadensis*) in Alberta, Canada, 2002–2019; (**B, E, H**) Mongolian gazelle (*Procapra gutturosa*) in Russian Transbaikalia, 2000–2020; and (**C, F, I**) wildebeest (*Connochaetes taurinus*) in the Serengeti-Mara ecosystem, 1961–2007. (**A-C**) In all three populations, individual migration probabilities vary substantially from year to year, such that a single migration probability does not accurately represent temporal dynamics. *Dashed lines* and *shaded regions* correspond to migration probabilities and standard errors estimated across each dataset in its entirety; *points* and *error bars* correspond to the means and the standard errors on the migration probabilities estimated for each year independently (though errors in C are so small that they are largely obscured by the points). (**D-F**) Total population size is highly predictive of the number of migrants but (**G-I**) has limited predictive power for explaining the number of residents. *Points* correspond to annual subpopulation size estimates, *error bars* reflect counting error. *Regression lines* correspond to the best fitting model (simplest model with ΔAIC_c_ < 2); a *solid line* represents a slope significantly different from 0, while a *dashed line* represents a slope of 0. See *Materials and Methods* for analytical details. Banff elk data (**A, D, G**) are from Martin *et al.* (2022), Mongolian gazelle data (**B, E, H**) are from Kirilyuk (2021), and Serengeti wildebeest data (**C, F, I**) are from Hopcraft *et al.* (2015). For elk panels, *dark blue lines/points* correspond to years through 2010, *light blue* to years after 2010, when there was an apparent state-change in the system. For gazelle panels, the *black outlined point* corresponds to the year 2007, an exceptionally mild winter during which nearly no Mongolian gazelle migrated (49). For wildebeest panels, *orange points* are the two outlier years immediately following the elimination of rinderpest, and the *orange line* is the best fitting model including these two outliers; *red points* are normal years, and the *red line* is the best fitting model without outliers. Note that panel axes are on different scales.

**FIGURE 2.**
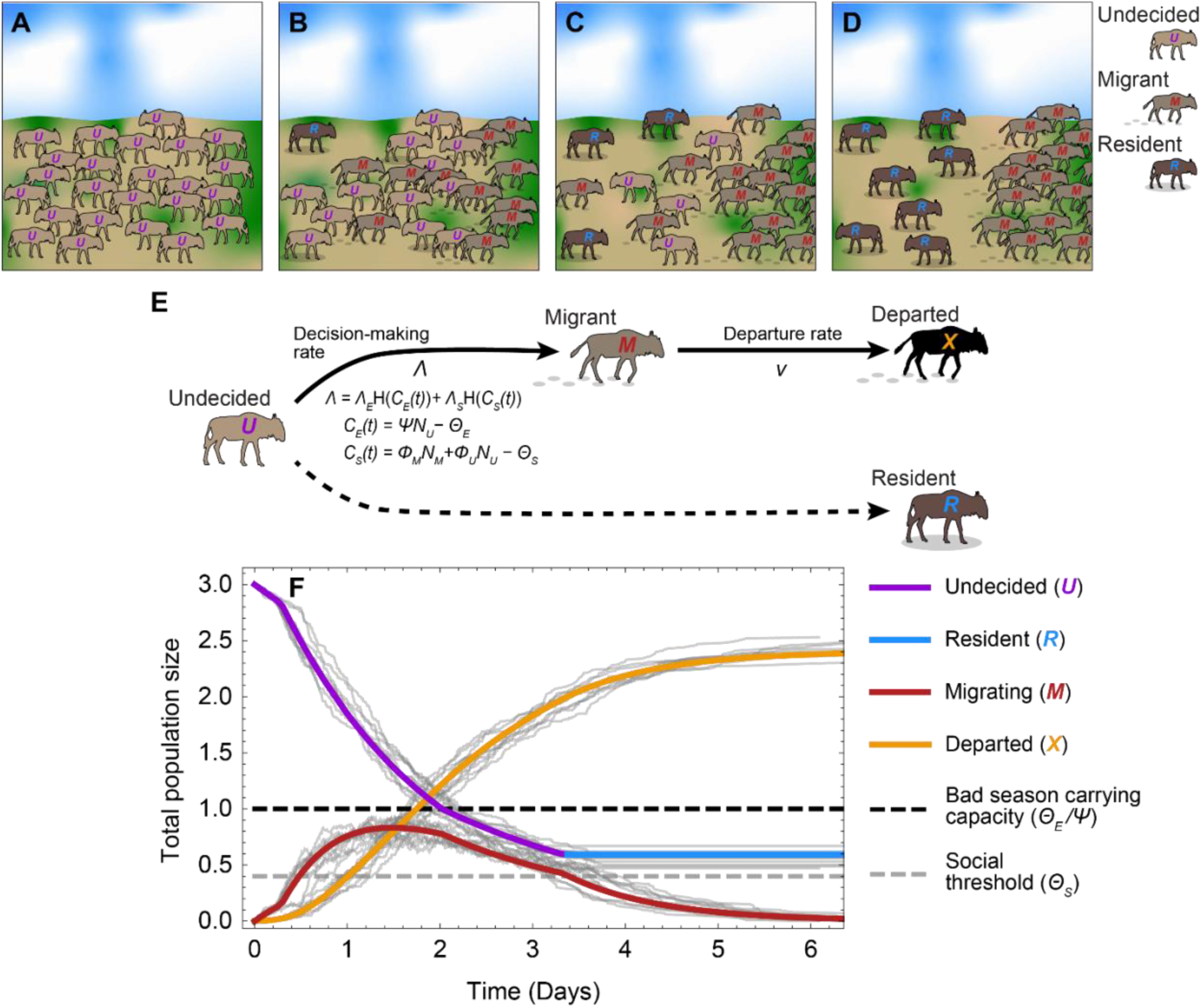
Schematic of model dynamics. We hypothesize that residents are undecided individuals that fail to start migrating before cues to migrate wane. We represent this visually in (**A-D**): (**A**) At high initial densities on the seasonal range, all individuals are undecided (*purple curve/symbols*) and therefore have the potential to become migratory. (**B**) As the good season gives way to the bad season, migratory cues exceed some threshold and some individuals stochastically decide to start migrating (*red curve/symbols*) at rate Λ. (**C**) Migrants leave the seasonal range at rate *v*, transitioning to the departed class (*orange curve/symbols*), and the intensity of migratory cues diminishes, with the result that some undecided individuals become fixed as residents (*light blue curve/symbols*) for the season. (**D**) At the end of the population partitioning process, migrating individuals have departed the seasonal range, and the remaining undecided individuals have all become residents. *Light gray curves* in (**F**) show ten independent realizations of the stochastic model, whereas the *thicker colored curves* are the result of integrating the deterministic model. Total population size is expressed relative to the bad season carrying capacity. Parameters: *U*_0_ = 3; *Θ*_S_ = 0.4; *ϕ*_M_ = 0.8; *ϕ*_U_ = 0.1; *Θ*_E_ = 1; *ψ* = 1; *ν* = 1; Λ_E_ = 0.4; Λ_S_ = 0.4.

## RESULTS

### Testing for a fixed migration probability

We first test whether individual migration probabilities vary on an annual basis or remain largely constant over several decades within the only three ungulate populations for which such data are available: Banff elk, Mongolian gazelle, and Serengeti wildebeest. For each population, we estimated a global migration probability across all years and its associated uncertainty. We then repeated this estimation for each year individually and compared the resulting yearly estimates to the global migration probability for each dataset.

Using this approach, we find that migration probabilities vary yearly in all three populations (**Fig. 1**). For the Banff elk, migration probabilities are highly variable when each year is modeled independently, and individual yearly migration probabilities overlap the global migration probability (the migration probability that best fits the entire dataset) for only 6 out of 18 years (*i.e.*, the 95% confidence intervals for both probabilities overlap; **Fig. 1A**). Even when accounting for uncertainty in the number of migrants and residents, which is substantial within the elk dataset, the estimated global migration probability is never able to explain more than half of the individual annual estimates (**Fig. S1**). For the Mongolian gazelle, the migration probability that best fits the entire dataset overlaps none of the individual years’ migration probabilities, which vary substantially, ranging from nearly 0 (*i.e.,* no migration, as in 2007) to nearly 1 (*i.e.,* complete migration) (**Fig. 1B**). Similarly, for the Serengeti wildebeest, the estimated global migration probability overlaps none of the individually estimated annual migration probabilities, which likewise vary substantially across years (**Fig. 1C**). In other words, year-to-year variation in migratory behavior within these three ungulate populations cannot be explained by a single, genetically prescribed migration probability, which supports the idea that an individual’s probability of migrating in a given year is shaped by local context.

### Empirical scaling of residents with total population size

Next, we quantify whether interannual variability in migration probability modifies the relationship between migrant/resident numbers and total population size (both expected to scale linearly and positively if migration probability were genetically fixed). To do so, we modeled the number of migrants and residents separately as functions of total population size and used model selection to identify the best-fitting relationship in each case. Within the three ungulate populations, we find that total population size, while highly predictive of migrant numbers (**Fig. 1D-F**), has limited predictive power for explaining variation in the number of residents (**Fig. 1G-I**).

Comparing linear models with and without population size as a predictor variable, we find that the best model of migrant Banff elk includes both population size and regime (pre- vs. post-2010, when an apparent system change occurred in Banff; see *Materials and Methods*) as predictors (**Table S1**); the number of migrants is strongly positively correlated with total population size in this model (slope = 0.984, *N* = 18, *t* = 15.170, *P* < 0.001; **Fig. 1D**). In contrast, the best model of resident Banff elk includes only population regime as a predictor (**Fig. 1G**).

For Mongolian gazelles, which underwent rapid range expansion during the study period, we model population density rather than total population size because the extent of their seasonal range varied substantially across years. The best model of migrant densities again includes total population density as a predictor (**Table S1**), and densities of migrants are highly correlated with overall population densities in this model (slope = 0.971, *N* = 21, *t* = 37.326, *P* < 0.001; **Fig. 1E**). For residents, the intercept-only model is the best model of resident population densities (**Table S1**), and resident densities are not significantly correlated with overall population densities when total population density is included as a predictor (slope = 0.029, *N* = 21, *t* = 1.100, *P* = 0.285; **Fig. 1H**).

Lastly, for the Serengeti wildebeest, the best models of migratory wildebeest numbers include total population size regardless of whether outlier years (the two years after the elimination of rinderpest from the Serengeti, when the system was likely still equilibrating) are included (**Table S1**); the number of migrants is strongly positively correlated to total population size both with (slope = 0.963, *N* = 27, *t* = 57.912, *P* < 0.001) and without (slope = 0.984, *N* = 25, *t* =110.127, *P* < 0.001) outlier years (**Fig. 1F**). Though the best model of resident wildebeest does include total population size as a predictor when outliers are included (**Table S1**), resident wildebeest numbers are only weakly correlated with total population size (slope = 0.037, *N* = 27, *t* = 2.241, *P* = 0.034; **Fig. 1I**). When outliers are excluded, the best model of resident numbers is the intercept-only model (**Table S1**), and residents are not significantly correlated to total population size when population size is included as a predictor (slope = 0.016, *N* = 25, *t* = 1.780, *P* = 0.088; **Fig. 1I**). As such, while the number of migrants clearly scales with total population size within these three ungulate populations, total population size has limited predictive power for explaining the number of residents—further evidence that the decision to migrate is not genetically fixed, but rather context-dependent.

### Model of migration onset

To investigate the context-dependence of the population-level partitioning into migrants and residents, we develop a model of migration onset wherein an individual’s decision to migrate is probabilistic and dependent on local environmental and/or social factors (4, 7, 13, 37–41). We use this model towards three aims: (i) to identify the conditions under which such a process results in partial migration (*i.e.,* the partitioning of the population into non-zero resident and migratory subpopulations); (ii) to evaluate the consequences of using social in addition to environmental information when deciding to migrate; and (iii) to explore potential drivers of year-to-year variability in the population-partitioning process.

The empirically established fact that observed character differences (*e.g.*, age, sex, body size) cannot account for within-population variation in ungulate migratory behavior suggests that the population-partitioning phenomenon itself, though potentially influenced quantitatively by character differences, is neither fully explained nor fully driven by such heterogeneities (12). Consequently, our initial model assumes a completely homogeneous population, *i.e.,* all individuals are assumed to be identical with respect to model parameters, to investigate whether spontaneous population partitioning into residents and migrants can occur nonetheless. Subsequently, we relax this assumption to explore how heterogeneities can further shape the emergent partitioning and contribute to the observed variability.

To begin, we assume that all individuals start in state *U*, *i.e.*, undecided about whether to migrate (**Fig. 2A**). The initial population size is equal to the growing (good) season carrying capacity, *U*_0_ (**Table 1**). Consistent with the fact that the migration onset is a rapid but not instantaneous event in many ungulate populations (4, 12, 26, 42), our model allows migration onset to be viewed as a temporally explicit process during which decisions are being made quickly (on the scale of hours to days), but not instantaneously, by individuals within a dynamic social context. As some undecided individuals begin to migrate, two additional states arise: migrating, *M*, individuals who have just started to migrate but have yet to leave the seasonal range (**Fig. 2B-C**), and departed, *X*, individuals who have already left the seasonal range. Since our model deals only with the onset of migration in one range and not with the processes that maintain the migration once it has begun, any departed individuals are assumed to be gone from the system (**Fig. 2**). Within our modeling framework, residents, *R*, are not a separate state; they are the outcome of the model: *i.e.,* they are the individuals that never transition out of the undecided state (**Fig. 2E**). Finally, because the onset of migration occurs over a short period (typically in a matter of days; 4, 12, 24, 40) during which demographic changes (births and deaths) are negligible, we assume that the total population size is constant, *i.e.*, *N*_U_(*t*) + *N*_M_(*t*) + *N*_X_(*t*) = *U*_0_ (where *N*_α_(*t*) is the number of individuals in class α={*U*, *M*, *X*} at time *t*). In other words, although individuals move between classes (*e.g.,* from resident to migrant to departed), these transitions represent redistribution within the population rather than demographic gains or losses.

**TABLE 1.**
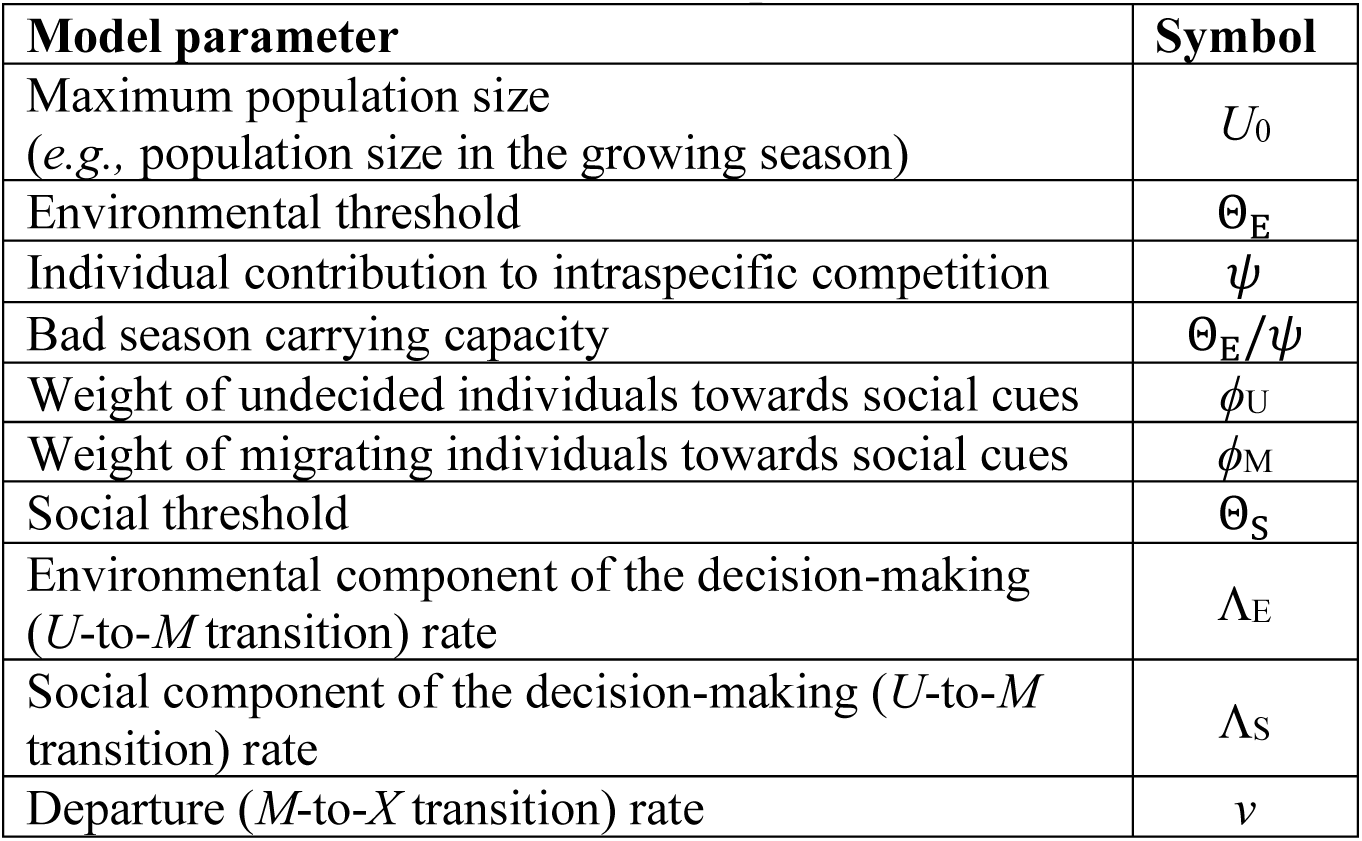
Model parameters.

Individuals decide to migrate (transition from undecided to migrating) probabilistically at rate Λ and depart the seasonal range (transition from migrating to departed) probabilistically at rate *v* (**Fig. 2E**, **Table 1**). These transition rates define the average time (on the scale of hours to days) that individuals spend in the undecided and migrating states, respectively. Therefore, *v* is a proxy for how long individuals remain in the seasonal range after starting to migrate, which implicitly introduces the effect of space in our model.

It is empirically well-established that environmental cues contribute to the decision to migrate (13, 21, 24, 42–44), but it is increasingly recognized that social information likely also contributes (16, 33, 34, 38). We therefore assume that the decision-making rate is controlled by the intensity of two classes of cues: environmental and social cues, *C*_E_ and *C*_S_ respectively, such that Λ = Λ(*C*_E_, *C*_S_). Our model is indifferent to the exact nature of these two cue classes, which remains broadly unclear empirically—they may be visual, auditory, olfactory, or pheromonal, or they might target some other sensory modality entirely (38–40).

In the case of environmental cues, competition appears to be a strong candidate for the ultimate mechanism by which cues are sensed and integrated by individuals (24, 35); for instance, removing forage competition (*e.g*., by providing supplemental forage; 45, 46) suppresses migratory behavior (2, 12, 23). We therefore assume that the intensity of the environmental cues, *C*_E_(*t*), is equal to the intensity of competition ‘sensed’ by each individual. We assume that only undecided individuals exert competitive pressure because migrating individuals are actively moving rather than foraging and should thus contribute minimally to local competition (this assumption can be easily relaxed and the results remain qualitatively robust). Accordingly, we define the intensity of the environmental cues perceived by a focal individual as the difference between the number of undecided individuals, *N*_U_(*t*), weighted by a nondimensional conversion factor *ψ* that accounts for per-capita resource demands, and that individual’s perception of resource availability in the environment, Θ_E_ (its ‘environmental threshold’):

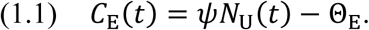

Because all individuals are initially undecided, the intensity of environmental cues decreases over time as individuals decide to migrate and depart (**Fig. 2**); the contribution of environmental cues to decision-making reaches zero when *ψN*_U_(*t*) = Θ_E_. Thus, Θ_E_/*ψ* emerges from our model as the bad season carrying capacity, *i.e.*, the maximum number of individuals the seasonal range can support during the bad season (**Table 1**).

While the role and nature of social cues in migratory decision-making is less certain, they likely do contribute to some degree, at least for certain ungulate populations (16, 33, 34, 38). In our model, we allow for the possibility that all sympatric individuals could contribute to social ‘signaling’, regardless of whether or not they begin to migrate. However, because migrating and undecided individuals might contribute differently to social cues (16, 33, 38, 41),we assign them different weights (*ϕ*_M_ and *ϕ*_U_, respectively; **Table 1**). We assume that the intensity of social cues, *C*_S_, is determined by the weighted combination of the number of undecided and migrating individuals relative to their social threshold, Θ_S_, which represents an individual’s sensitivity to social influence:

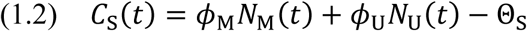

Finally, we assume that individuals integrate environmental and social cues differently when deciding whether to migrate (39, 40), and that the two cue types thus contribute independently to the decision-making rate. Hence, Λ is a linear combination of two step-like response functions:

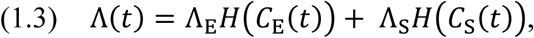

where *H* is the step function: *H*(*x*) = 1 if *x* > 0 and *H*(*x*) = 0 otherwise. *C*_E_ and *C*_S_ are given by Eqs. (1.1) and (1.2), respectively, and Λ_E_ and Λ_S_ are the environmental and social components of the decision-making rate, respectively (**Table 1**). Choosing step response functions simplifies the mathematical treatment of our model because the decision-making rate remains a linear function of the number of individuals in each class. From Eq. (1.3), Λ can take four different values—specifically, Λ = {0, Λ_E_, Λ_S_, Λ_E_ + Λ_S_}—depending on the intensity of the social and environmental cues. We constrain our analyses to cases where Λ_E_ ≠ 0, as it is well documented that environmental cues always play a role in migration (13, 21, 24, 42–44). Thus, though a potential parameterization of the model, we do not consider the case where Λ = Λ_S_ (*i.e.*, where only social cues determine ungulate migratory decision-making) due to its biological implausibility.

Because transitions between states are probabilistic, the state of the whole population at any point in time is defined by the distribution of individuals across their possible internal states *U*, *M*, and *X*. Since this probability distribution function cannot be obtained analytically, we can perform numerical simulations of the stochastic dynamics (*thin gray curves* in **Fig. 2F**) or use standard techniques to derive a deterministic model approximation. This approximation consists of a system of coupled ordinary differential equations describing how the mean number of individuals within each state changes with time (47) (see **Supplementary Material** for detailed derivations):

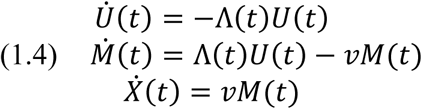

where the dots indicate time derivatives. The departure rate *v* is constant, while the decision-making rate Λ depends on time because it is a function of environmental and social cues, per Eq. (1.3). Finally, note that the last equation is redundant, as we assumed a constant total population size for the duration of the population-partitioning event. **Figure 2F** confirms that the deterministic approximation accurately captures the mean temporal dynamics of the system (see **Fig. S2** in the **Supplementary Materials** for a more detailed comparison).

### Theoretical results

Because our empirical analyses show that the context-dependency of migratory decision-making directly impacts how the numbers of migrants and residents scale with total population size (**Fig. 1**), and because these subpopulations are the empirical observable, we present our theoretical results in terms of these subpopulation sizes to facilitate the generation of testable predictions.

#### Residents are a robust phenomenon in our model, but their numbers are largely not predictable from total population size

We find that, for a broad parameter regime, our model can reproduce a population partitioning process that results in a non-zero resident population (**Fig. 2E**). We also find that the number of migrants increases near-linearly with increasing total population across the entire range of population sizes within our model (**Fig. 3A**), while the trends in the resident population are more nuanced (**Fig. 3B**).

**FIGURE 3.**
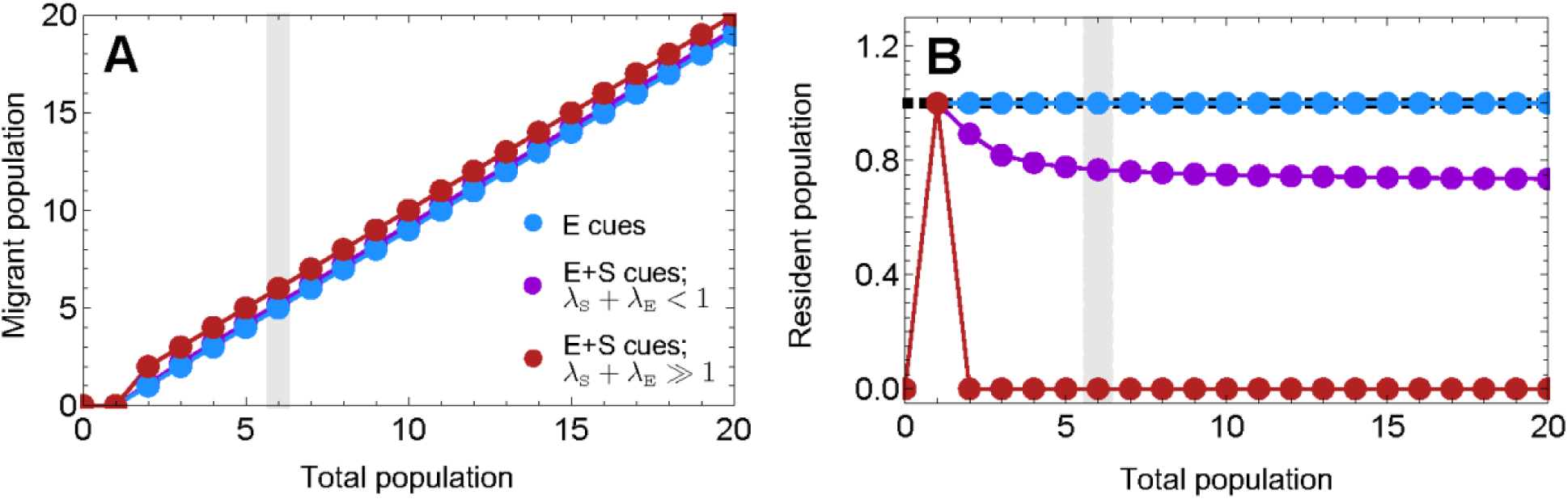
Model predictions for how the numbers of (A) migrants and (B) residents vary with total population size. The model recapitulates the empirical pattern that, (**A**) while the number of migrants is strongly positively correlated with total population size, (**B**) the number of residents is not necessarily predictable from total population size. Instead, the number of residents depends on the nature and strength of two types of cues: if only environmental cues drive migration (*blue curves*), the number of residents will plateau at the bad season carrying capacity (*black dashed line*); if social cues also contribute to decision-making and the decision-making rate is slower than the departure rate (*purple curves*), the number of residents will be lower than the bad season carrying capacity (except at small population sizes, where social cues are weak and thus the number of residents equals the bad season carrying capacity); otherwise, when social cues contribute to decision-making and the decision to migrate is made very fast relative to the rate at which migrants leave (*red curves*), then residents can entirely disappear. The *grey shaded region* corresponds to the total population size used in figure 4. Subpopulation sizes are expressed relative to the bad season carrying capacity. Parameters: *Θ*_S_ = 0.4; *ϕ*_M_ = 0.8; *ϕ*_U_ = 0.1; *Θ*_E_ = 1; *ψ* = 1; *ν* = 1; (Λ_E_, Λ_S_) = (0.4, 0.2) *purple*; (0.4, 0.0) *blue*; (15, 15) *red*.

Specifically, we find that the relationship between the number of residents and total population size depends on: (i) the total population size and (ii) the different sources of cues triggering migration (**Table 1**). At small initial population sizes, regardless of the type of cues, all individuals remain resident because per-capita resources are plentiful and competition is minimal, and any social impetus to migrate is weak due to the small number of conspecifics (**Fig. 3B**). As a result, the number of residents scales positively with total population size at this stage, reflecting normal population growth; in such cases, the population would not yet be considered partially migratory, but rather a growing or establishing population. Once the population is large enough for any migration to occur, the outcome of the population partitioning process depends strongly on whether migratory decision-making is driven solely by environmental cues or is also influenced by social cues. If migration is driven only by environmental cues (*blue curve* in **Fig. 3B**), then the number of residents plateaus at the bad season carrying capacity, *R*_*E*_ = Θ_*E*_/*ψ*. If social cues contribute to decision-making, we distinguish two different population-size regimes for how the number of residents scales with total population size. In the first, at intermediate total population sizes, the number of residents declines with population size (**Fig. 3B**). In the second, at sufficiently large population sizes, the resident population is either zero (*i.e.*, all individuals migrate; *red curve* in **Fig. 3B**) or it plateaus at a size that is independent of initial population size (*purple curve* in **Fig. 3B**). The model therefore robustly recapitulates the empirical pattern that the number of residents—but not the number of migrants—can be largely invariant with total population size (**Fig. 1**).

Within this latter regime (where the number of residents plateaus with population size), the existence of the resident population and its size both depend on the strength of social cues. When social cues contribute to migration, the two characteristic time scales of the model—the average time to make the decision to migrate, 1/(Λ_S_ + Λ_E_), and the average time for a migrant to leave the seasonal range, 1/*ν*—control a phase separation between complete migration (when Λ_S_ + Λ_E_ > *ν*) and partial migration (when Λ_S_ + Λ_E_ < *ν*). In other words, residents exist only if migrating individuals depart the seasonal range fast enough to leave behind those sparse, undecided individuals that have not started migrating (due to the probabilistic nature of the decision-making process). Thus, if social cues contribute to migratory decision-making, residents should exist only when departure rates are very rapid, faster than the rate at which individuals make decisions. When migration is partial (*i.e.*, Λ_S_ + Λ_E_ < *ν*), then the resident subpopulation, *R*_E+S_, satisfies

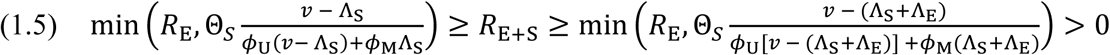

In other words, *R*_E+S_ can never exceed the bad-season carrying capacity *R*_*E*_ = Θ_*E*_/*ψ*, though it can be lower if *R*_E_ is greater than the fractions in (1.5) (see **Supplementary Material Section 1.4.3** for proof). Thus, social cues amplify environmental cues, making it more likely that migration is complete and reducing the number of residents relative to environmental cues alone.

#### Depending on the cues that determine migration, there are different sources of year-to-year variability in resident numbers

If migration is only driven by environmental cues, the number of residents is always equal to the bad season carrying capacity. Year-to-year variation in resident numbers thus arises predominantly from variation in the severity of the bad season, Θ_*E*_/*ψ* in our model, which can vary considerably in practice (13, 24, 48, 49). If social cues contribute to the decision to migrate, then additional drivers of variation in the number of residents could be: (i) the total population size, if it fluctuates within small-to-intermediate ranges; (ii) the social weights of migrating and undecided individuals (*ϕ*_M_, *ϕ*_U_; **Fig. 4A-B**); or (iii) the social threshold (Θ_*S*_; **Fig. 4C**). Indeed, all these parameters likely do vary between populations and for a particular population from year to year as environmental and/or social contexts change, which could generate observed variability in resident numbers (**Fig. 1**). For example, as newly established populations accumulate more information about their environments, migrating individuals may carry more weight in the decision-making process (increasing *ϕ*_M_) (38, 41), causing the number of residents to decrease over successive migratory seasons, as has been observed empirically (16).

**FIGURE 4.**
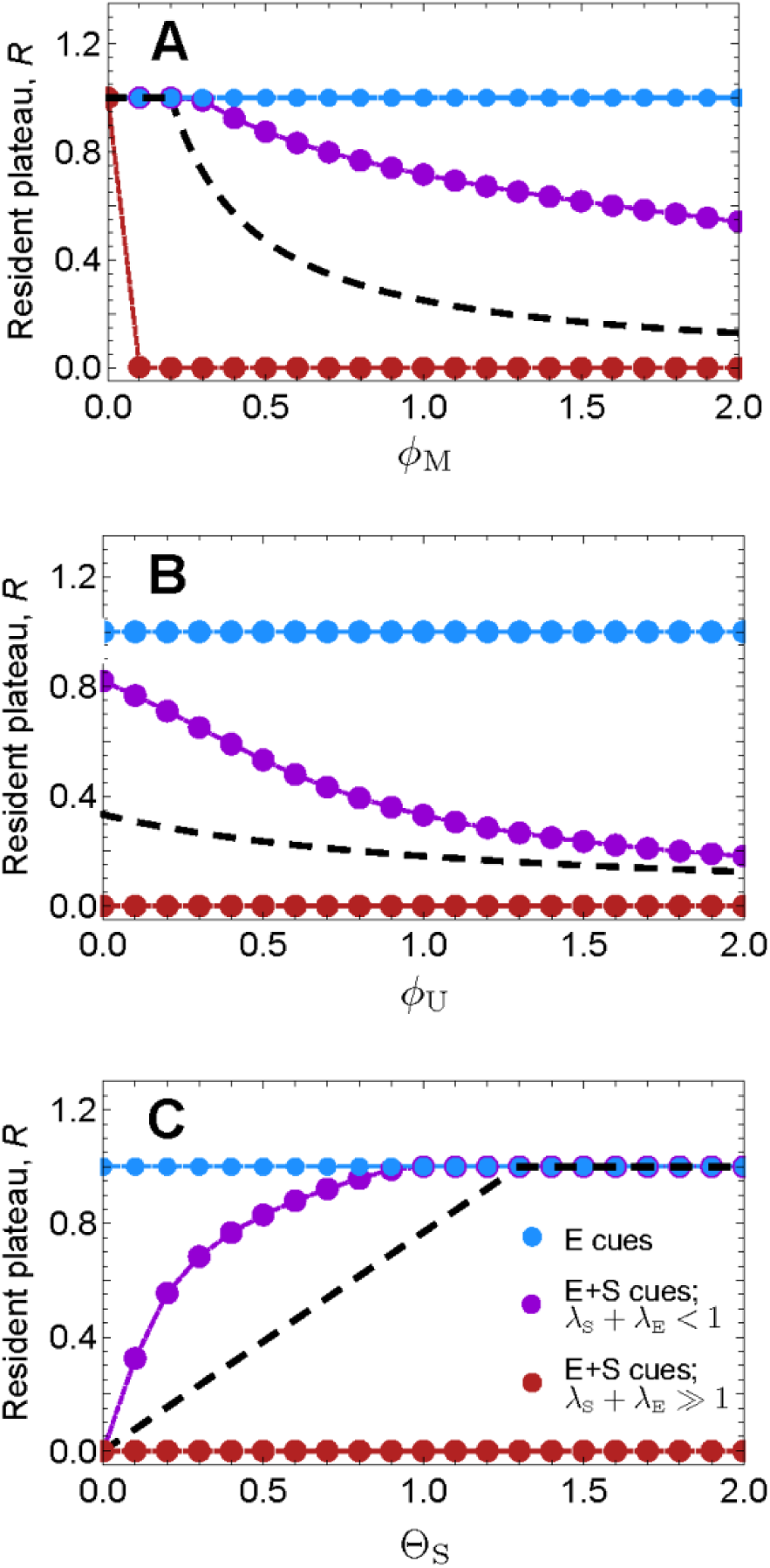
The dependence of the resident plateau on (A) the contribution of migrating individuals to social cues, (B) the contribution of undecided individuals to social cues, and (C) the social threshold. Parameters related to social cues have no effect on the resident plateau when social cues do not contribute to the decision to migrate (*blue curves*) or when the decision to migrate is made very fast relative to the rate at which migrants leave (*red curves*). Between these extremes, when social cues contribute to migratory decision-making (*purple curves*), the resident plateau declines as (**A**) migrating and (**B**) undecided individuals contribute more to social cues and as (**C**) the social threshold declines. The *black dashed curve* is the analytical estimate within the lower bound provided in Eq. (1.5). Parameters (unless used as the x-axis control parameter): U_0_ = 6; *Θ*_S_ = 0.4; *ϕ*_M_ = 0.8; *ϕ*_U_ = 0.1; *Θ*_E_ = 1; *ψ* = 1; *ν* = 1; (Λ_E_, Λ_S_) = (0.4, 0.2) *purple*; (0.4, 0.0) *blue*; (15, 15) *red*.

#### Character differences within the population can further impact resident numbers

All results above were derived assuming that, within one year, there are no differences between individuals with respect to the model parameters. Although the homogeneity constraint was instrumental to explore the source of the population-partitioning process, our model can readily incorporate character differences to investigate how heterogeneities shape the emergent resident plateau and impact year-to-year variation. To illustrate this, we apply our model to a simple scenario in which the population is comprised of two classes of individuals: small- and large-bodied. For simplicity, we only introduce heterogeneity in thresholds and in the parameters that determine the weights of undecided and migrating individuals, but the same approach can be extended to consider heterogeneities in any parameters. We make the biologically plausible assumption that small individuals contribute less to environmental cues, *ψ*_s_ < *ψ*_ℓ_, due to lower per capita resource demands. For the same reason, we assume that small individuals are less environmentally sensitive and thus have higher environmental thresholds, Θ_Es_ > Θ_Eℓ_ (though we also consider the scenario where Θ_Es_ = Θ_Eℓ_; **Fig. S3**). Finally, we assume that, when social cues contribute to migratory decision-making, larger individuals contribute more strongly to social cues (resulting from their social dominance and/or greater cultural knowledge; 16, 33, 34), such that large individuals have higher *ϕ*_U_ and *ϕ*_M_ than small individuals. We do not *a priori* specify a relationship between the social thresholds of the two size classes, Θ_Ss_ and Θ_Sℓ_, instead exploring how different relationships between the social thresholds impact patterns. We compare the two scenarios that yield partial migration (*i.e.,* a non-zero resident population): (1) migration driven solely by environmental cues, in the parameter regime where Θ_*E*_/*ψ* > 0; and (2) migration driven by both environmental and social cues, in cases where Λ_S_ + Λ_E_ < *ν*.

We ask two questions: first, does heterogeneity impact the magnitude of the resident plateau, implying that natural year-to-year variation in the relative abundance of the two body-size classes could contribute to variation in resident numbers? And second, is there a skew in the contribution of the two body-size classes to the resident population relative to the initial population, as is widely observed in empirical partially migratory populations (12, 22–27)?

Under our simple assumptions, the answers to both questions are yes; these qualitative patterns persist even when the environmental thresholds of the two size classes are assumed to be equal (**Fig. S3**). First, the magnitude of the resident plateau is influenced by heterogeneity, with higher initial fractions of the smaller, less environmentally sensitive individuals resulting in more residents (**Fig. 5A**). Second, there is always skewed representation of the two classes in the resident population relative to the total population, but the direction of the skew can vary (**Fig. 5B**). If migration is determined only by environmental cues, then small individuals are overrepresented in the resident population, regardless of their prevalence in the initial population. If social cues also contribute to migratory decision-making, the skew depends on the relative magnitudes of the social thresholds of the two classes. If small individuals have higher social thresholds than large ones (*i.e.*, Θ_Ss_ > Θ_Sℓ_), then the skew is in the same direction as before: small individuals remain overrepresented in the resident population. However, if social thresholds have the opposite relationship to environmental ones (*i.e.,* Θ_Ss_ < Θ_Sℓ_), then large individuals can become overrepresented among residents when they are abundant in the initial population (as in **Fig. 5B**). These results can be reframed as size class-specific probabilities of migrating (**Fig. 5C-D**): for example, when migration is determined solely by environmental cues, small individuals always have a lower probability of migrating than large individuals, and their migration probability declines as the number of small individuals in the initial population increases.

**FIGURE 5.**
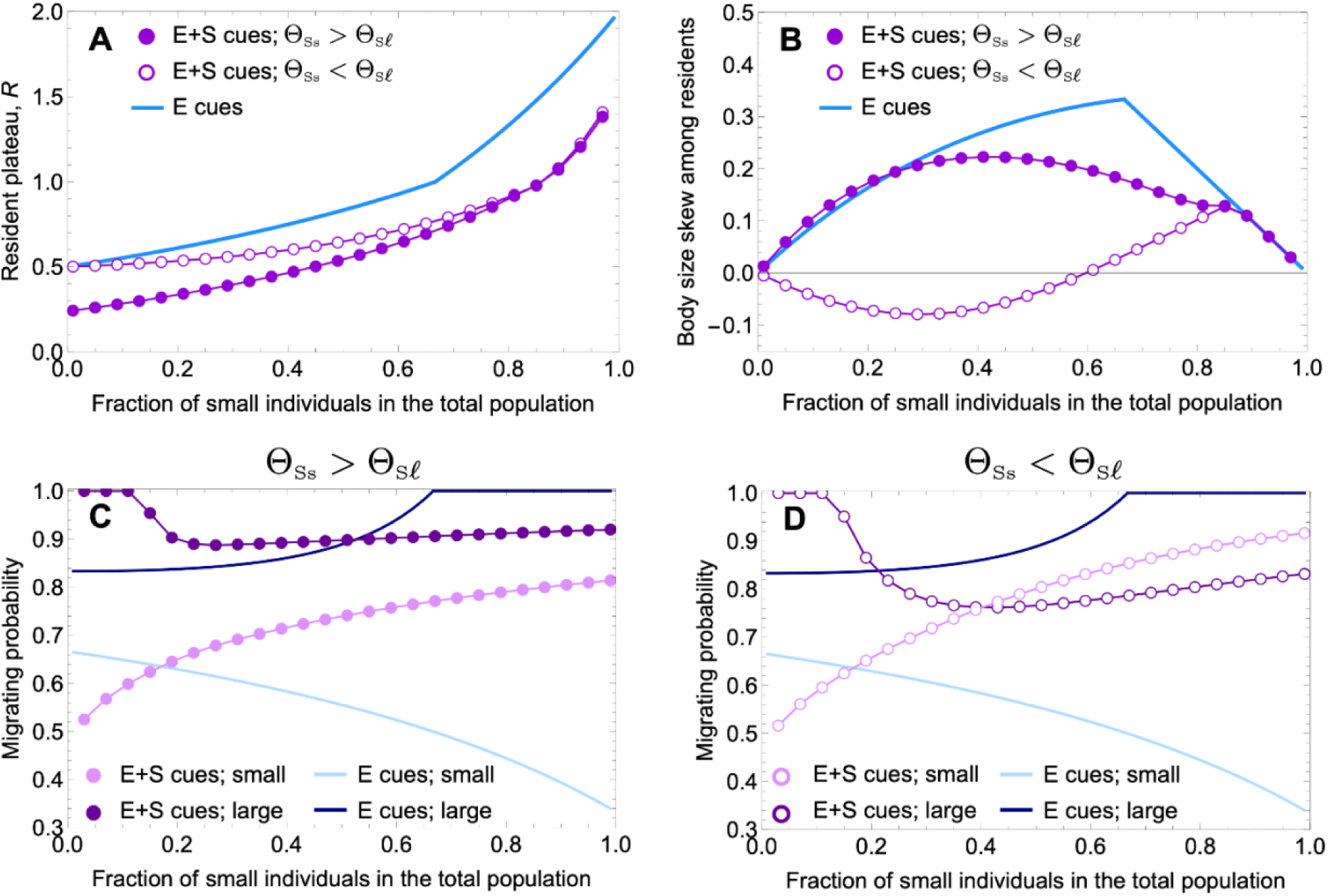
Model predictions for heterogeneous populations. (**A**) The resident plateau also depends on the proportion of small and large-bodied individuals in the population, as well as on the relationship between their social thresholds. (**B**) If social cues do not factor into the decision to migrate, the proportion of small individuals is always higher in the resident subpopulation than in the total population. When social cues contribute, the overrepresentation of small individuals in the resident subpopulation remains when Θ_Ss_ > Θ_Sℓ_; but when Θ_Ss_ < Θ_Sℓ_ , this bias is frequency-dependent and small individuals can be underrepresented in the resident subpopulation if they are infrequent in the overall population. (**C, D**) These skews within the resident subpopulation translate into probabilities that individuals from each size class migrate: (**C**) small individuals are always less likely to migrate than large individuals when Θ_Ss_ > Θ_Sℓ_, but (**D**) these probabilities become frequency-dependent when Θ_Ss_ < Θ_Sℓ_. Parameters: *U*_0_ = 3; *ϕ*_Mℓ_ = 0.8; *ϕ*_Ms_ = 0.4; *ϕ*_Uℓ_ = 0.1; *ϕ*_Us_ = 0.08; Λ_S_ = 0.4; Λ_E_ = 0.4; *ψ*_ℓ_ = 1; *ψ*_s_ = 0.5; *ν* = 1; Θ_Es_ = 1; Θ_Eℓ_ = 0.5; Θ_Ss_ = 0.4; Θ_Sℓ_ = 0.8 or Θ_Sℓ_ = 0.2. See **Supplementary Material** for definitions of bias metrics and migration probabilities.

## DISCUSSION

Here we explore how ungulates decide whether or not to migrate each year. We observe that migration probabilities are highly variable across years within three different ungulate populations and that, though the number of migrants scales positively with total population size, the number of residents does not. These empirical results suggest that the decision to migrate is context-dependent. We therefore present and analyze a model of ungulate migration in which the decision to migrate is probabilistic and dependent on local cues: individuals probabilistically start migrating in accordance with the intensity of environmental and/or social cues. Our model allows migration onset to be viewed as a temporally explicit process during which decisions are made quickly (on the scale of hours to days) but not instantaneously by individuals within a dynamic context. This is consistent with the fact that the onset of migration is a rapid but not instantaneous event in many ungulate populations (4, 12, 26, 42). We find that this individual decision-making robustly produces residents for a broad parameter regime, conforming to empirical observations that residents exist within many migratory ungulate populations (12) and theoretical expectations that both tactics can be fitness-maximizing under certain conditions (18, 19, 36). Our model predicts that such a non-zero resident subpopulation is *mechanistically achievable* only when, during the fast migration onset, migrating individuals depart their seasonal range faster than the average rate at which individuals make the decision to migrate.

When only environmental cues factor into migratory decision-making, the size of the resident subpopulation is set by the bad season carrying capacity. When social cues also contribute to decision-making, the two cue classes amplify each other twofold: they narrow the parameter regime where partial migration occurs and, when partial migration does occur, they lower the number of residents left behind. Thus, migration driven by both social and environmental cues produces fewer residents and is more likely to produce no residents at all. The size of the resident subpopulation resulting from our model scales variably with total population size. At small initial population sizes, the number of residents increases simply because the population is growing—migration has yet to begin. At intermediate population sizes, the number of residents can decrease if social cues contribute to migratory decision-making. At sufficiently large population sizes, the number of residents plateaus and is no longer predictably related to total population size, which aligns with empirical patterns observed in several migratory ungulate populations. In contrast, the number of migrants *does* scale positively with total population size in our model, also consistent with empirical observations.

Our model further sheds light on what—if not population size—might underlie the observed variation in resident numbers: in addition to some (small) variability resulting from the fact that the decision to migrate is inherently probabilistic, there can also be year-to-year variation in other parameters that control the resident plateau, such as the environmental threshold, the social threshold, and the contributions of migrating and undecided individuals towards social cues. For example, as ungulates accumulate information about a landscape and migratory routes are established, the contributions of migrating and/or undecided individuals to social cues might increase (38, 40, 41), causing a population to become increasingly migratory as experience accrues over successive migratory seasons (16). In particular, our model recapitulates empirical observations that especially strong environmental cues, such as those occurring during drought (50), can cause complete migration within partially migratory populations or trigger some migration in populations that usually do not migrate (27, 50, 51). Within our modeling framework, such a scenario might correspond to a decrease in the environmental threshold and reduction in the bad season carrying capacity. On the flip side, when environmental cues are weak, such as when there are minimal seasonal differences in carrying capacity, our model predicts that migration can entirely collapse. This accords with empirical findings that migratory behavior can be suppressed when the bad season is not severe—such as in 2007 in Mongolia when the bad season was abnormally mild and almost no Mongolian gazelle migrated (see 49)—or when seasonal differences in carrying capacity are artificially removed, such as by providing supplemental forage (45, 46). As such, interannual changes in the environmental or social contexts of a population likely contribute to observable variation in the number of residents through time.

That our model can account for within-population variation in the number of residents supports our assumption that, within ungulates, the decision to migrate depends in part upon the strength of local cues. However, our results simultaneously reveal that there is more involved in the decision to migrate than just local cues. For instance, our model predicts a low probability of the same individual staying resident multiple years in a row when residents are rare. Yet this result is inconsistent with observations that resident individuals can remain resident for multiple years, even when residents are rare (12, 19, 27). This discrepancy between our model results and empirical observations implies that the decision to migrate may be a multi-year process (19, 52), a suggestion that is consistent with mounting evidence that ungulates learn when, where, and whether to migrate through prior experience (both one’s own and others’; 15, 16, 33, 34, 53, 54). While the model we present here only considers the onset of a single migratory season (and so cannot account for learning or for fitness consequences), extensions of this modeling framework to consider multi-year migratory dynamics are feasible and can, *e.g.*, incorporate switching lags, whereby there is some inertia to switching migratory tactics across years. Similarly, empirical evidence suggests that dynamics in parts of the landscape beyond the seasonal range can impact migratory decision-making: for instance, high densities in adjacent ranges can act as ‘social fences’, inhibiting out-migration (12, 55, 56). Thus, multi-patch extensions of this modeling framework might couple decision-making across patches, incorporating this additional source of information as well.

All results discussed above arose in the absence of any character-based heterogeneities in the initial population that would bias towards or against migration. Nonetheless, our model reproduces a resident subpopulation, suggesting that the population partitioning process does not necessitate heterogeneities. This finding accords with empirical observations that the same individual will switch between migratory tactics across years (12), as well as with the lack of consistent genetic and character differences between migrants and residents (12, 29–34). That said, though inconsistent across populations, character and demographic differences between residents and migrants are widely observed in partially migratory populations: migrant and resident subpopulations have been shown to differ in average body size, sex ratios, age distributions, and pregnancy rates (12, 22–27). If such differences do translate into differences in migratory propensity between individuals within a population, as has long been theorized (12, 57, 58), then character-dependent migration propensities could quantitatively alter the partitioning process. Using body size as an example, we demonstrate that heterogeneity in thresholds between size classes can both lower the number of residents and skew size distributions within the resident population relative to the initial population. However, the direction of this skew is not *a priori* predictable: we find that small individuals are more likely to remain resident than large individuals in the absence of social cues, but social cues can invert this pattern, making it more likely for small individuals to migrate when they are rare to begin with. The frequency-dependence of this results cautions against attempting to extract simple behavioral rules from observed character skews: in our example, if there are more than expected small individuals among residents, it does not necessarily imply that small individuals have higher social thresholds than large individuals; they could actually have *lower* social thresholds and be abundant within the population. Future research is therefore needed to determine how observed character differences map to heterogeneities in migratory propensity across diverse ungulate populations, and the consequent patterns for migrants and residents.

Beyond ungulates, our results may have broader implications for understanding the drivers and fitness consequences of collective behaviors across biological systems. Partial migration has also been documented in insects, fish, and birds (20, 44, 55, 59). While these taxa are subject to different constraints and often migrate for different reasons (*e.g.,* for breeding purposes; 20, 21, 40, 44, 60), similar context-dependent decision-making processes could underlie their migrations. For instance, in some fish and bird systems, it has been hypothesized that resource acquisition relative to a genetically influenced threshold can determine whether individuals remain resident or migrate, a process conceptually similar to that captured by our model (12, 21, 44). And beyond migration, the emergence of diverse behavioral states from imperfectly coordinated collective behavior has been observed in a variety of systems, including flowering synchronization (61) and slime mold aggregations (62, 63). Research in some of these systems suggests that imperfect synchronization may both be a general progenitor of behavioral diversity and also contribute to the robustness of the whole population. For example, imperfect cell-to-cell coordination during slime mold aggregation leads to a similar population partitioning event between a multicellular and unicellular phase (62, 64, 65), which could serve as a bet-hedging strategy that favors population survival in uncertain environmental conditions (62, 64, 65). The same could be true for migratory ungulates: the existence of residents may be critical for ensuring population resilience in the face of unpredictable climatic conditions, which is particularly relevant in light of global change and an increasingly unpredictable biosphere. All in all, stochastic decision-making informed by local cues is likely a broad phenomenon, and one that may have evolutionary and ecological implications for collective systems. Further research on the potential for, and mechanisms underlying, behavioral diversification across varied biological systems is thus key to identifying general principles and fitness consequences of collective behaviors.

## MATERIAL AND METHODS

### Empirical data on migration probability

To quantify empirical migration probabilities within ungulate populations, we searched Web of Science for longitudinal counts of resident and migrant individuals within a population. Our search terms were migra* AND residen* AND (population size OR population density OR total population OR population count OR census) AND (annual OR yearly OR longitudinal) AND (herbivore OR mammal OR ungulate OR Bov* OR Cerv* OR Equ*); we supplemented the results of this search with our own knowledge of the literature. This process yielded 86 papers (as of February 2022), but only three included data from partially migrating ungulate populations: Banff elk (*Cervus canadensis;* 19), Mongolian gazelle (*Procapra gutturosa;* 66) and Serengeti wildebeest (*Connochaetes taurinus*; 26). We extracted the data from these publications using WebPlotDigitizer.

All three species are gregarious, sexually dimorphic ungulates in the order Artiodactyla, with males larger than females. They live in fission-fusion societies, forming large mixed-sex groups led by dominant males as well as smaller affiliate groups of subdominant bachelor males (26, 67). All are also polygynous and breed seasonally once per year, with females generally giving birth to a single offspring at a time (26, 67). Mongolian gazelle are the smallest of the three species, at *ca.* 28 kg (5, 67), and are sometimes described as nomadic, lacking well-established migratory routes (49, 54, 66). Elk and wildebeest are larger, at 130 and 180 kg respectively (5, 67), and tend to have consistent migratory routes that they follow year after year (2, 19, 26).

The Banff elk dataset spans 18 years (2002–2019) and consists of minimum estimates of migrant and resident numbers based on winter aerial survey counts (19, 68); counting uncertainty within the dataset is substantial. The dataset spans an apparent system change that occurred in 2010, perhaps related to the extirpation of a sympatric population of caribou (*Rangifer tarandus*) from the region (69), the canonization of an additional migratory route (19), and/or the intensification of predation pressure within their migration corridor (70). The Mongolian gazelle dataset covers 21 years (2000–2020) with ground transect counts conducted 2–3 times per year, alongside seasonal range size measurements; no explicit counting uncertainties are reported. These data capture a period of rapid territory expansion, during which the size of the seasonal range increased dramatically (66), as well as environmental extremes, including the unusually mild winter of 2007 when nearly no individuals migrated (49). The Serengeti wildebeest dataset includes 27 years of data spanning several decades (1961–2007), compiled from multiple sources and methods, including ground and aerial surveys; counting uncertainties are reported for only some years (26). These data span the elimination of rinderpest from the Serengeti-Mara ecosystem, during which time the population rapidly expanded (26). The two years following the elimination of rinderpest were characterized by anomalously high resident numbers, possibly the result of population dynamics re-equilibrating.

### Testing for a fixed migration probability

For each of these three datasets, we used a two-step statistical analysis, implemented in Python3 (71), to evaluate whether empirical data on residents and migrants through time could result from a distribution with constant migration probability *p*: if each individual in a population migrates with probability *p* and stays as a resident with complementary probability 1-*p*, then the decision-making process can be modeled as a Bernoulli trial, and the number of migrants, *M*, in a population with *N* individuals follows a binomial distribution,

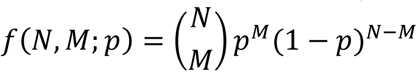

So, we first merged all years of data for a given species and used maximum likelihood estimation to fit the data to a binomial distribution. This provided an estimate of the probability of migrating, *p̂*_*T*_, that best fits the entire dataset. For a binomial distribution, we can maximize the log-likelihood manually and obtain the value of *p̂*_*T*_ analytically

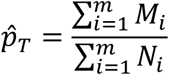

where *M*_*i*_ and *N*_*i*_ are the number of migrants and the total population size, respectively, for each year *i* in the dataset. We calculated 95% confidence intervals around the global migration probabilities per Clopper and Pearson (1934), using ‘binomtest’ in SciPy (73) (**Fig. 1A-C**).

In the second step of the analysis, we tested whether each year of data could be generated from a binomial distribution centered on this migration probability *p̂*_*T*_ for each population. We again used maximum likelihood estimation, but this time we fitted separate binomial distributions to each year of data independently. This analysis returns *m* different estimates of the migration probability *p̂*_*i*_ = *M*_*i*_⁄*N*_*i*_ for *i* = 1,…, *m* along with their corresponding 95% confidence intervals, again obtained using the function ‘binomtest’ from SciPy (73). We then assessed whether the annual migration probabilities *p̂*_*i*_ were significantly different from the global migration probability *p̂*_*T*_ for each population (*i.e.*, the 95% confidence intervals of the two probabilities did not overlap).

Lastly, to account for the substantial counting uncertainty on the number of residents and migrants in the elk dataset, we conducted a sensitivity analysis using the two-step statistical approach described above. From reported counting errors, we generated 10^5^ datasets consisting of *m_elk_=*18 pairs of resident-migrant subpopulation sizes: we generated each pair of data points by sampling a random number from a Normal distribution centered at the reported resident and migrant subpopulation sizes, with variability given by the counting errors reported in (19). For each of these randomized datasets, we obtained *p̂*_*T*_ and the year-specific *p̂*_*i*_s, and we quantified the fraction of years for which the migration probability *p̂*_*i*_ differed significantly from the dataset-wide migration probability *p̂*_*T*_. Finally, we merged these values for all the 10^5^ randomized datasets and constructed a histogram of the fraction of years that could be explained by the overall migration probability (**Fig. S1**).

### Empirical scaling of residents with total population size

We further examined how resident and migrant numbers scaled with total population size for each of the three populations. Using model selection in R v3.6.1 (74), we fitted linear models to each dataset that did and did not (*i.e.,* intercept-only) include total population size as a predictor (**Table S1**). We modeled the numbers of residents and migrants separately to capture potential differences in their scaling relationships and to avoid obscuring non-linear patterns that might arise given context-dependent migratory decision-making.

Each dataset required unique modeling decisions due to the unique ecological contexts of each. For the Banff elk, we split the dataset into two population regimes, pre- and post-2010, to account for the system change that seemingly occurred around 2010; we included regime as an additional predictor in model selection, which substantially improved model fit (**Table S1**). For the Mongolian gazelle, because the size of their seasonal range changed dramatically during the study period, we converted resident and migrant subpopulation totals into subpopulation densities, either directly from measured seasonal range area or from range area estimates generated by linearly interpolating from measured values (**Figs 1E, 1H**). Finally, for the Serengeti wildebeest, we constructed separate model sets with and without the two post-rinderpest outlier years to evaluate the influence of these outliers on scaling relationships (**Fig. 1F, 1I; Table S1).**

We then compared model-corrected Akaike Information Criterion (AIC_c_) to assess model fit (**Table S1**). We consider all models with ΔAIC_c_ < 2 but consider the simplest model within ΔAIC_c_ < 2 as the overall ‘best’ model for the purposes of interpretation (see also 5).

## ACKNOWLEDGEMENTS

We would like to thank Thomas A. Morrison for illuminating email exchanges about the Serengeti wildebeest migration that inspired this model. We would like to gratefully acknowledge Rob Pringle, Merlijn Staps, Justin M. Calabrese, Luisa Ramirez, Dan Rubenstein, Jonathan Levine, and the Pringle and Tarnita Lab Groups at Princeton University for their valuable feedback on this project. JOA was supported by the NSF Graduate Research Fellowship Program (Fellow ID: 2019256075). RMG is partially supported by the Simons Foundation through grant 284558FY19; FAPESP through the BIOTA Jovem Pesquisador grant 2019/05523-8 and ICTP-SAIFR grant 2021/14335-0. This work is partially funded by the Center of Advanced Systems Understanding (CASUS) which is financed by Germany’s Federal Ministry of Education and Research (BMBF) and by the Saxon Ministry for Science, Culture and Tourism (SMWK) with tax funds on the basis of the budget approved by the Saxon State Parliament.

## DATA ACCESSIBILITY STATEMENT

Data are available in the supplementary materials and in Dryad Data Repository (DOI: 10.5061/dryad.n02v6wx77).

## Notes

### Competing Interest Statement

The authors have declared no competing interest.

## REFERENCES

1. A. P. Dobson, et al., Road will ruin Serengeti. Nature 467, 272–273 (2010).

2. T. M. Anderson, et al., Interplay of competition and facilitation in grazing succession by migrant Serengeti herbivores. Science 383, 782–788 (2024).

3. F. Larsen, et al., Wildebeest migration drives tourism demand in the Serengeti. Biological Conservation 248, 108688 (2020).

4. M. J. Kauffman, et al., Causes, Consequences, and Conservation of Ungulate Migration. Annual Review of Ecology, Evolution, and Systematics 52, null (2021).

5. J. O. Abraham, G. P. Hempson, J. T. Faith, A. C. Staver, Seasonal strategies differ between tropical and extratropical herbivores. Journal of Animal Ecology 91, 681–692 (2022).

6. J. M. Fryxell, J. Greever, A. R. E. Sinclair, Why are Migratory Ungulates So Abundant? The American Naturalist 131, 781–798 (1988).

7. S. D. Albon, R. Langvatn, Plant Phenology and the Benefits of Migration in a Temperate Ungulate. Oikos 65, 502–513 (1992).

8. A. C. Staver, G. P. Hempson, Seasonal dietary changes increase the abundances of savanna herbivore species. Science Advances 6, eabd2848 (2020).

9. S. Bauer, B. Hoye, Migratory Animals Couple Biodiversity and Ecosystem Functioning Worldwide. Science 344, 1242552 (2014).

10. D. T. Bolger, W. D. Newmark, T. A. Morrison, D. F. Doak, The need for integrative approaches to understand and conserve migratory ungulates. Ecology Letters 11, 63–77 (2008).

11. G. Harris, S. Thirgood, J. G. C. Hopcraft, J. P. G. M. Cromsigt, J. Berger, Global decline in aggregated migrations of large terrestrial mammals. Endangered Species Research 7, 55–76 (2009).

12. J. E. Berg, M. Hebblewhite, C. C. St. Clair, E. H. Merrill, Prevalence and Mechanisms of Partial Migration in Ungulates. Front. Ecol. Evol. 7, 325 (2019).

13. E. O. Aikens, et al., Wave-like Patterns of Plant Phenology Determine Ungulate Movement Tactics. Current Biology 30, 3444–3449.e4 (2020).

14. J. O. Abraham, N. S. Upham, A. Damian-Serrano, B. R. Jesmer, Evolutionary causes and consequences of ungulate migration. Nat Ecol Evol 1–9 (2022). 10.1038/s41559-022-01749-4.

15. C. Bracis, T. Mueller, Memory, not just perception, plays an important role in terrestrial mammalian migration. Proceedings of the Royal Society B: Biological Sciences 284, 20170449 (2017).

16. B. R. Jesmer, et al., Is ungulate migration culturally transmitted? Evidence of social learning from translocated animals. Science 361, 1023–1025 (2018).

17. J. M. Fryxell, A. R. E. Sinclair, Causes and consequences of migration by large herbivores. Trends in Ecology & Evolution 3, 237–241 (1988).

18. B. van Moorter, et al., Seasonal release from competition explains partial migration in European moose. Oikos 130, 1548–1561 (2021).

19. H. W. Martin, M. Hebblewhite, E. H. Merrill, Large herbivores in a partially migratory population search for the ideal free home. Ecology 103 (2022).

20. P. Lundberg, The evolution of partial migration in Birds. Trends Ecol Evol 3, 172–175 (1988).

21. B. B. Chapman, C. Brönmark, J.-Å. Nilsson, L.-A. Hansson, The ecology and evolution of partial migration. Oikos 120, 1764–1775 (2011).

22. M. Festa-Bianchet, Seasonal range selection in bighorn sheep: conflicts between forage quality, forage quantity, and predator avoidance. Oecologia 75, 580–586 (1988).

23. P. J. White, T. L. Davis, K. K. Barnowe-Meyer, R. L. Crabtree, R. A. Garrott, Partial migration and philopatry of Yellowstone pronghorn. Biological Conservation 135, 502–510 (2007).

24. N. J. Singh, L. Börger, H. Dettki, N. Bunnefeld, G. Ericsson, From migration to nomadism: movement variability in a northern ungulate across its latitudinal range. Ecological Applications 22, 2007–2020 (2012).

25. A. M. Fudickar, A. Schmidt, M. Hau, M. Quetting, J. Partecke, Female-biased obligate strategies in a partially migratory population. Journal of Animal Ecology 82, 863–871 (2013).

26. J. G. C. Hopcraft, et al., “6. Why Are Wildebeest the Most Abundant Herbivore in the Serengeti Ecosystem?” in Serengeti IV: Sustaining Biodiversity in a Coupled Human-Natural System, (University of Chicago Press, 2015), pp. 125–174.

27. S. L. Eggeman, M. Hebblewhite, H. Bohm, J. Whittington, E. H. Merrill, Behavioural flexibility in migratory behaviour in a long-lived large herbivore. Journal of Animal Ecology 85, 785–797 (2016).

28. A. Kaitala, V. Kaitala, P. Lundberg, A Theory of Partial Migration. The American Naturalist 142, 59–81 (1993).

29. N. J. Georgiadis, “Population Structure of Wildebeest: Implications for Conservation” in *Serengeti II: Dynamics*, Management, and Conservation of an Ecosystem, (1995).

30. D. W. Coltman, J. G. Pilkington, J. M. Pemberton, Fine-scale genetic structure in a free-living ungulate population. Molecular Ecology 12, 733–742 (2003).

31. A. D. McDevitt, et al., Survival in the Rockies of an endangered hybrid swarm from diverged caribou (Rangifer tarandus) lineages. Molecular Ecology 18, 665–679 (2009).

32. E. M. Ernest, et al., “The Serengeti Migration: Genetic differentiation of migratory and non-migratory wildebeest” in Norwegian School of Veterinary Science, (2012).

33. K. K. Barnowe-Meyer, P. J. White, L. P. Waits, J. A. Byers, Social and genetic structure associated with migration in pronghorn. Biological Conservation 168, 108–115 (2013).

34. K. E. Colson, K. S. White, K. J. Hundertmark, Parturition site selection in moose (Alces alces): evidence for social structure. Journal of Mammalogy 97, 788–797 (2016).

35. J. A. Stabach, “Movement, resource selection, and the physiological stress response of white-bearded wildebeest,” Colorado State University, United States -- Colorado. (2015).

36. B. van Moorter, et al., Consequences of barriers and changing seasonality on population dynamics and harvest of migratory ungulates. Theor Ecol 13, 595–605 (2020).

37. R. M. Holdo, R. D. Holt, J. M. Fryxell, Opposing Rainfall and Plant Nutritional Gradients Best Explain the Wildebeest Migration in the Serengeti. The American Naturalist 173, 431–445 (2009).

38. W. K. Oestreich, et al., The influence of social cues on timing of animal migrations. Nat Ecol Evol 6, 1617–1625 (2022).

39. D. W. Winkler, et al., Cues, strategies, and outcomes: how migrating vertebrates track environmental change. Mov Ecol 2, 10 (2014).

40. S. Bauer, et al., “Cues and decision rules in animal migration” in Animal Migration: A Synthesis, (Oxford University Press, 2011), pp. 68–87.

41. I. D. Couzin, J. Krause, N. R. Franks, S. A. Levin, Effective leadership and decision-making in animal groups on the move. Nature 433, 513–516 (2005).

42. M. P. Laforge, M. Bonar, E. Vander Wal, Tracking snowmelt to jump the green wave: phenological drivers of migration in a northern ungulate. Ecology 102 (2021).

43. E. O. Aikens, et al., The greenscape shapes surfing of resource waves in a large migratory herbivore. Ecol Lett 20, 741–750 (2017).

44. A. Ferguson, T. E. Reed, T. F. Cross, P. McGinnity, P. A. Prodöhl, Anadromy, potamodromy and residency in brown trout Salmo trutta: the role of genes and the environment. Journal of Fish Biology 95, 692–718 (2019).

45. J. D. Jones, et al., Supplemental feeding alters migration of a temperate ungulate. Ecological Applications 24, 1769–1779 (2014).

46. K. Barker, Home Is Where the Food Is: Causes and Consequences of Partial Migration in Elk. *Graduate Student Theses, Dissertations*, & Professional Papers (2018).

47. R. Toral, P. Colet, Stochastic Numerical Methods: An Introduction for Students and Scientists (John Wiley & Sons, 2014).

48. M. Valeix, H. Fritz, S. Chamaillé-Jammes, M. Bourgarel, F. Murindagomo, Fluctuations in abundance of large herbivore populations: insights into the influence of dry season rainfall and elephant numbers from long-term data. Animal Conservation 11, 391–400 (2008).

49. K. A. Olson, et al., A mega-herd of more than 200,000 Mongolian gazelles Procapra gutturosa: a consequence of habitat quality. Oryx 43, 149–153 (2009).

50. J. O. Abraham, G. P. Hempson, A. C. Staver, Drought-response strategies of savanna herbivores. Ecology and Evolution 9, 7047–7056 (2019).

51. M. E. Nelson, Winter range arrival and departure of white-tailed deer in northeastern Minnesota. Can. J. Zool. 73, 1069–1076 (1995).

52. T. A. Morrison, D. T. Bolger, Wet season range fidelity in a tropical migratory ungulate. Journal of Animal Ecology 81, 543–552 (2012).

53. T. Boulinier, E. Danchin, The use of conspecific reproductive success for breeding patch selection in terrestrial migratory species. Evol Ecol 11, 505–517 (1997).

54. R. Martínez-García, J. M. Calabrese, T. Mueller, K. A. Olson, C. López, Optimizing the Search for Resources by Sharing Information: Mongolian Gazelles as a Case Study. Phys. Rev. Lett. 110, 248106 (2013).

55. E. Matthysen, Density-dependent dispersal in birds and mammals. Ecography 28, 403–416 (2005).

56. A. Mysterud, et al., Partial migration in expanding red deer populations at northern latitudes – a role for density dependence? Oikos 120, 1817–1825 (2011).

57. T. H. Clutton-Brock, Reproductive Effort and Terminal Investment in Iteroparous Animals. The American Naturalist 123, 212–229 (1984).

58. F. B. Bercovitch, C. P. Loomis, R. G. Rieches, Age-Specific Changes in Reproductive Effort and Terminal Investment in Female Nile Lechwe. Journal of Mammalogy 90, 40–46 (2009).

59. J. Cote, et al., Behavioural synchronization of large-scale animal movements - disperse alone, but migrate together?: Synchronization of large-scale animal movements. Biol Rev 92, 1275–1296 (2017).

60. A. K. Shaw, Drivers of animal migration and implications in changing environments. Evol Ecol 30, 991–1007 (2016).

61. D. H. Janzen, Why Bamboos Wait So Long to Flower. Annual Review of Ecology, Evolution, and Systematics 7, 347–391 (1976).

62. C. E. Tarnita, A. Washburne, R. Martinez-Garcia, A. E. Sgro, S. A. Levin, Fitness tradeoffs between spores and nonaggregating cells can explain the coexistence of diverse genotypes in cellular slime molds. PNAS 112, 2776–2781 (2015).

63. F. W. Rossine, R. Martinez-Garcia, A. E. Sgro, T. Gregor, C. E. Tarnita, Eco-evolutionary significance of “loners.” PLoS Biol 18, e3000642 (2020).

64. D. Dubravcic, M. van Baalen, C. Nizak, An evolutionarily significant unicellular strategy in response to starvation in Dictyostelium social amoebae. F1000Res 3, 133 (2014).

65. R. Martínez-García, C. E. Tarnita, Seasonality can induce coexistence of multiple bet-hedging strategies in *Dictyostelium discoideum* via storage effect. Journal of Theoretical Biology 426, 104–116 (2017).

66. V. E. Kirilyuk, Rapidly increasing migratory activity of Mongolian gazelle (Procapra gutturosa) and the sightings of Goitered gazelle (Gazella subgutturosa) in Transbaikalia as an alarm. Rus.J.Theriol. 20, 25–30 (2021).

67. H. Wilman, et al., EltonTraits 1.0: Species-level foraging attributes of the world’s birds and mammals. Ecology 95, 2027–2027 (2014).

68. M. Hebblewhite, et al., Is the Migratory Behavior of Montane Elk Herds in Peril? The Case of Alberta’s Ya Ha Tinda Elk Herd. Wildlife Society Bulletin 34, 1280–1294 (2006).

69. M. Hebblewhite, C. White, M. Musiani, Revisiting Extinction in National Parks: Mountain Caribou in Banff. Conservation Biology 24, 341–344 (2010).

70. M. Hebblewhite, E. H. Merrill, Demographic balancing of migrant and resident elk in a partially migratory population through forage–predation tradeoffs. Oikos 120, 1860–1870 (2011).

71. G. Van Rossum, F. Drake, Python 3. (2009). Deposited 2009.

72. C. J. Clopper, E. S. Pearson, The Use of Confidence or Fiducial Limits Illustrated in the Case of the Binomial. Biometrika 26, 404–413 (1934).

73. P. Virtanen, et al., SciPy 1.0: fundamental algorithms for scientific computing in Python. Nat Methods 17, 261–272 (2020).

74. R Core Team, R: A language and environment for statistical computing. (2023). Deposited 2023.

