## Supplemental Text 1 for "Should I stay or should I go? Modelling the decision-making process behind ungulate partial migration"

### Supplementary Material for: 'Should I stay or should I go? Modeling the decision-making process behind ungulate partial migration'

Joel O. Abraham*[1], Ricardo Martinez-Garcia*[,2,3,4], Finote Gijsman†[1], Erin M. Phillips§[1], and Corina E. Tarnita[1]

[1]Department of Ecology and Evolutionary Biology, Princeton University, Princeton, New Jersey, USA
[2]Center for Advanced Systems Understanding (CASUS); Helmholtz-Zentrum Dresden-Rossendorf (HZDR), Görlitz, Germany
[3]ICTP South American Institute for Fundamental Research and Instituto de Física Teórica, Universidade Estadual Paulista - UNESP, São Paulo, Brazil
[4]Department of Ecology, Institute of Biosciences, University of São Paulo, São Paulo, Brazil

**3 Model supplementary figures** 16

*These authors contributed equally to this work and share first authorship
†These authors contributed equally to this work

### Part I

### Empirical data analysis

| Population | | | Predictors | $R^2$ | df | logLik | $AIC_c$ | $\Delta AIC_c$ | weight |
|---|---|---|---|---|---|---|---|---|---|
| Ya Ha Tinda elk | | Resident | ***regime*** | ***0.939*** | ***3*** | ***-86.26*** | ***180.23*** | ***0.000*** | ***0.686*** |
| Ya Ha Tinda elk | | Resident | regime*total_population | 0.953 | 5 | -83.94 | 182.89 | 2.657 | 0.182 |
| Ya Ha Tinda elk | | Resident | regime+total_population | 0.939 | 4 | -86.22 | 183.52 | 3.292 | 0.132 |
| Ya Ha Tinda elk | | Resident | total_population | 0.668 | 3 | -101.51 | 210.74 | 30.511 | 0.000 |
| Ya Ha Tinda elk | | Resident | single intercept only | 0.000 | 2 | -111.44 | 227.67 | 47.443 | 0.000 |
| Ya Ha Tinda elk | | Migrant | *regime*total_population* | *0.963* | *5* | *-83.94* | *182.89* | *0.000* | *0.579* |
| Ya Ha Tinda elk | | Migrant | ***regime+total_population*** | ***0.952*** | ***4*** | ***-86.22*** | ***183.52*** | ***0.635*** | ***0.421*** |
| Ya Ha Tinda elk | | Migrant | total_population | 0.736 | 3 | -101.51 | 210.74 | 27.854 | 0.000 |
| Ya Ha Tinda elk | | Migrant | regime | 0.212 | 3 | -111.37 | 230.45 | 47.559 | 0.000 |
| Ya Ha Tinda elk | | Migrant | single intercept only | 0.000 | 2 | -113.51 | 231.82 | 48.934 | 0.000 |
| Mongolian gazelle | | Resident | ***intercept only*** | ***0.000*** | ***2*** | ***16.48*** | ***-28.30*** | ***0.000*** | ***0.674*** |
| Mongolian gazelle | | Resident | *total_population* | *0.060* | *3* | *17.13* | *-26.85* | *1.449* | *0.326* |
| Mongolian gazelle | | Migrant | ***total_population*** | ***0.987*** | ***3*** | ***17.13*** | ***-26.85*** | ***0.000*** | ***1.000*** |
| Mongolian gazelle | | Migrant | intercept only | 0.000 | 2 | -28.11 | 60.88 | 87.733 | 0.000 |
| Serengeti wildebeest | With outliers | Resident | ***total_population*** | ***0.167*** | ***3*** | ***-315.83*** | ***638.71*** | ***0.000*** | ***0.768*** |
| Serengeti wildebeest | With outliers | Resident | intercept only | 0.000 | 2 | -318.30 | 641.10 | 2.398 | 0.232 |
| Serengeti wildebeest | With outliers | Migrant | ***total_population*** | ***0.993*** | ***3*** | ***-315.83*** | ***638.71*** | ***0.000*** | ***1.000*** |
| Serengeti wildebeest | With outliers | Migrant | intercept only | 0.000 | 2 | -382.07 | 768.64 | 129.93 | 0.000 |
| Serengeti wildebeest | Without outliers | Resident | *total_population* | *0.121* | *3* | *-275.75* | *558.64* | *0.000* | *0.578* |
| Serengeti wildebeest | Without outliers | Resident | ***intercept only*** | ***0.000*** | ***2*** | ***-277.36*** | ***559.27*** | ***0.630*** | ***0.422*** |
| Serengeti wildebeest | Without outliers | Migrant | ***total_population*** | ***0.998*** | ***3*** | ***-275.75*** | ***558.64*** | ***0.000*** | ***1.000*** |
| Serengeti wildebeest | Without outliers | Migrant | intercept only | 0.000 | 2 | -354.12 | 712.79 | 154.14 | 0.000 |

**Table S1:** Model selection results for the predictors of resident and migrant subpopulation size for different ungulate populations. 'Preferred' models (all models with ΔAICc < 2) are *italicized*, whereas the 'best' model (simplest model with ΔAICc < 2) is **bolded**.

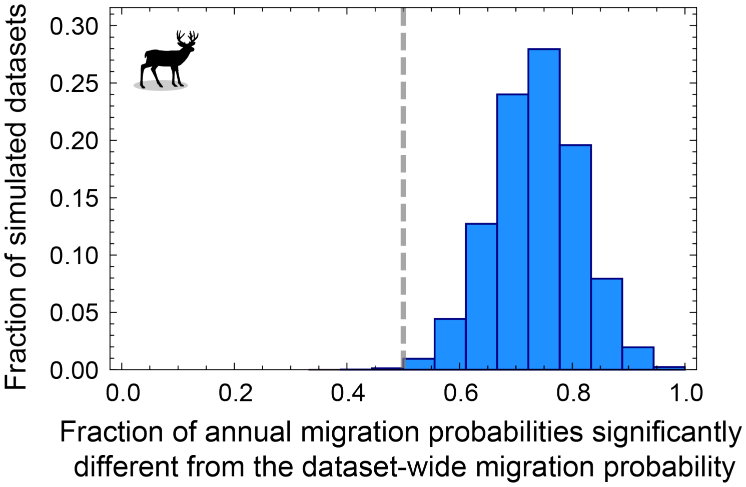

**Fig. S1:** Because counting uncertainty within the elk dataset is substantial, we generated $10^5$ datasets consisting of 18 pairs (one for each year of data) of resident-migrant subpopulation sizes: we generated each pair of data points by sampling a random number from a Normal distribution centered at the reported resident and migrant population sizes, with variability given by the corresponding reported counting errors. For each of these randomized datasets, we obtained a dataset-wide migration probability and the year-specific migration probabilities; we then quantified the fraction of years for which the year-specific migration probabilities differed significantly from the dataset-wide migration probability for all the $10^5$ randomized datasets. A single dataset-wide migration probability was never able to explain more than half of the individual annual estimates. The dashed gray line indicates 0.5.

#### Part II

### Model analysis

#### 1 Monomorphic population

##### 1.1 Stochastic model formulation

We introduce a spatially implicit stochastic modeling framework to explore whether we can recapitulate empirical patterns of partial migration across three ungulate species. In our model, individuals are identical and can be in one of three possible states: undecided $U$, migrating $M$, and departed $X$. We assume that all individuals are initially in the undecided state and that the total population size is equal to the seasonal-range carrying capacity. Mathematically, this initial condition means that $N_{\mathrm{M}}(0) = N_{\mathrm{X}}(0) = 0$ and $N_{\mathrm{U}}(0) = U_0$, where $U_0$ is the seasonal-range carrying capacity. Finally, because the typical time scales of migration onset are much smaller than demographic time scales, the model further assumes that the total population size is constant, $N_{\mathrm{U}}(t) + N_{\mathrm{M}}(t) + N_{\mathrm{X}}(t) = U_0$ for all $t$.

The dynamics consist of a series of individual transitions between $U$ and $M$, at time-dependent rate $\Lambda(t)$, and between $M$ and $X$, at constant rate $v$. Each of these rates gives the probability per unit time of the transition to occur and is thus non-negative. Reverse transitions are not allowed. Therefore, migrating individuals cannot become undecided, and departed individuals leave the seasonal range and cannot return to the migrating state. Using conventional notation, we can write these transitions in terms of biological reactions,

$$U \xrightarrow{\Lambda(t)} M \tag{1.1.1}$$

$$M \xrightarrow{v} X \tag{1.1.2}$$

The transition rate $v$ defines the typical time an individual stays in the range while in the migrating state and thus provides a proxy for the velocity at which migrating animals leave the seasonal range. $\Lambda(t)$ depends on the intensity of environmental and social cues, $C_{\mathrm{E}}$ and $C_{\mathrm{S}}$ respectively, through two step-like threshold response functions. We model these response functions using $\Theta$-Heaviside functions, such that $H(x) = 1$ for $x > 0$ and $H(x) = 0$ otherwise. This choice is mathematically convenient because it keeps the model linear in the number of individuals. We additionally assume that individuals process and integrate social and environmental cues independently, which leads to an $U$-to-$M$ transition rate of the form

$$\Lambda(t) = \Lambda_{\mathrm{E}} H(C_{\mathrm{E}}(t)) + \Lambda_{\mathrm{S}} H(C_{\mathrm{S}}(t)), \tag{1.1.3}$$

where the first term corresponds to the response to environmental cues, and the second one to the response to social cues.

As introduced in the main text, we assume that social cues are given by the difference between a weighted combination of the number of both undecided and migrating individuals and individual's sensitivity to social influence

$$C_{\mathrm{S}}(t) = \phi_{\mathrm{U}} N_{\mathrm{U}}(t) + \phi_{\mathrm{M}} N_{\mathrm{M}}(t) - \Theta_{\mathrm{S}}. \tag{1.1.4}$$

Environmental cues, as perceived by a focal undecided individual, are given by the difference between the number of undecided individuals, weighted by their *per-capita* resource demands, and the focal undecided individual's perception of resource availability (environmental threshold),

$$C_{\mathrm{E}}(t) = \psi N_{\mathrm{U}}(t) - \Theta_{\mathrm{E}}. \tag{1.1.5}$$

Because all individuals are initially in the undecided state, the intensity of environmental cues decreases over time and the environmental-cue contribution to $\Lambda(t)$ reaches zero when $\psi N_{\mathrm{U}}(t) = \Theta_{\mathrm{E}}$. Therefore, $\Theta_{\mathrm{E}}/\psi$ is the bad-season carrying capacity.

Given the probabilistic transitions between individual internal states in Eqs. (1.1.1) and (1.1.2), the distribution of individuals through the possible internal states, $P(N_{\mathrm{U}}, N_{\mathrm{M}}; t)$, defines the state of the ungulate population. Notice that because we assume that the total population size remains constant during the entire onset of migration, $N_{\mathrm{X}}(t)$ is a redundant variable. $P(N_{\mathrm{U}}, N_{\mathrm{M}}; t)$ changes with time according to

$$\begin{aligned} P(N_{\mathrm{U}}, N_{\mathrm{M}}, t+dt) &= \Lambda(t)(N_{\mathrm{U}}+1)P(N_{\mathrm{U}}+1, N_{\mathrm{M}}-1, t)dt + && (1.1.6)\\ & \quad v\,(N_{\mathrm{M}}+1)P(N_{\mathrm{U}}, N_{\mathrm{M}}+1, t)dt + && (1.1.7)\\ & \quad \Big[1-\Lambda(t)N_{\mathrm{U}}dt - vN_{\mathrm{M}}dt\Big]P(N_{\mathrm{U}}, N_{\mathrm{M}}, t) && (1.1.8)\end{aligned}$$

where $\Lambda(t)$ depends on time through the values of $N_{\mathrm{U}}$ and $N_{\mathrm{M}}$ and thus remains constant between $t$ and $dt$. In the limit $dt \to 0$, we obtain the Master equation for $P(N_{\mathrm{U}}, N_{\mathrm{M}}, t)$, a differential equation that describes the deterministic dynamics of the probabilistic distribution of individuals in the possible internal states (Toral and Colet, 2014; Van Kampen, 1992),

$$\begin{aligned} \frac{\partial P(N_{\mathrm{U}}, N_{\mathrm{M}}, t)}{\partial t} &= \Lambda(t)(N_{\mathrm{U}}+1)P(N_{\mathrm{U}}+1, N_{\mathrm{M}}-1, t) + \\ & \quad v(N_{\mathrm{M}}+1)P(N_{\mathrm{U}}, N_{\mathrm{M}}+1, t) - \Big[\Lambda(t)N_{\mathrm{U}} + vN_{\mathrm{M}}\Big]P(N_{\mathrm{U}}, N_{\mathrm{M}}, t). \quad (1.1.9)\end{aligned}$$

We cannot solve the Master equation (1.1.9) analytically because it is nonlinear through the dependence of $\Lambda$ on the number of both undecided and migrating individuals, but we can perform numerical simulations using standard methods like the Gillespie algorithm (Gillespie, 1977; Toral and Colet, 2014) (gray trajectories in main text Fig. 2) or do some approximations to obtain partial information about $P(N_{\mathrm{U}}, N_{\mathrm{M}}; t)$ through its moments.

#### 1.2 Deterministic approximation to the stochastic model

To gain analytical insights into the stochastic dynamics described by the Master equation (1.1.9), we reduce it to a set of coupled ordinary differential equations (ODEs) for the mean values of the number of individuals in each state (Toral and Colet, 2014). Here we show how to obtain the ODE for the mean number of undecided individuals, $U(t)$,

$$U(t) = \sum_{N_{\mathrm{U}}=0}^{\infty}\sum_{N_{\mathrm{M}}=0}^{\infty} N_{\mathrm{U}} P(N_{\mathrm{U}}, N_{\mathrm{M}}, t). \quad (1.2.1)$$

We can derive the equation for the mean number of migrating individuals following the same steps, whereas the equation for the mean number of departed individuals follows from the conservation of the total population size. Multiplying both sides of Eq. (1.1.9) by $N_{\mathrm{U}}$, summing over $N_{\mathrm{U}}$ and $N_{\mathrm{M}}$, and switching the order of the time derivative and the sums on the left side, we get

$$\begin{aligned} \frac{\partial}{\partial t}\sum_{N_{\mathrm{U}}=0}^{\infty}\sum_{N_{\mathrm{M}}=0}^{\infty} N_{\mathrm{U}} P(N_{\mathrm{U}}, N_{\mathrm{M}}, t) &= \sum_{N_{\mathrm{U}}=0}^{\infty}\sum_{N_{\mathrm{M}}=0}^{\infty} \Lambda(t)\, N_{\mathrm{U}}(N_{\mathrm{U}}+1)P(N_{\mathrm{U}}+1, N_{\mathrm{M}}-1, t) + \\ & \quad \sum_{N_{\mathrm{U}}=0}^{\infty}\sum_{N_{\mathrm{M}}=0}^{\infty} v\, N_{\mathrm{U}}(N_{\mathrm{M}}+1)P(N_{\mathrm{U}}, N_{\mathrm{M}}+1, t) - \\ & \quad \sum_{N_{\mathrm{U}}=0}^{\infty}\sum_{N_{\mathrm{M}}=0}^{\infty} N_{\mathrm{U}}\Big[\Lambda(t)\, N_{\mathrm{U}} + v\, N_{\mathrm{M}}\Big]P(N_{\mathrm{U}}, N_{\mathrm{M}}, t) \end{aligned} \quad (1.2.2)$$

Computing the sums on the left side of Eq. (1.2.2) and the third term of the right side already results in the mean number of undecided individuals. However, we need to do some manipulations in the indexes for the first and second terms on the right side. Defining a new pair of indexes $j \equiv N_{\text{M}} - 1$ and $i \equiv N_{\text{U}} + 1$, the first term on the right side of Eq. (1.2.2) becomes

$$\sum_{N_{\text{U}}=0}^{\infty} \sum_{N_{\text{M}}=0}^{\infty} \Lambda(t)\, N_{\text{U}}(N_{\text{U}}+1) P(N_{\text{U}}+1, N_{\text{M}}-1, t) = \sum_{i=1}^{\infty} \sum_{j=-1}^{\infty} \Lambda(t)(i-1)\, i\, P(i,j,t) \tag{1.2.3}$$

Next, because the terms corresponding to $i = 0$ and $j = -1$ are always zero, we can rewrite the range of the sums on the right side of Eq. (1.2.3) so they both start at zero. Finally, because $i$ and $j$ are indexes we can rename them $j \equiv N_{\text{M}}$ and $i \equiv N_{\text{U}}$ to get,

$$\sum_{i=1}^{\infty} \sum_{j=-1}^{\infty} \Lambda(t)\,(i-1)\, i\, P(i,j,t) = \sum_{N_{\text{U}}=0}^{\infty} \sum_{N_{\text{M}}=0}^{\infty} \Lambda(t)(N_{\text{U}}-1) N_{\text{U}} P(N_{\text{U}}, N_{\text{M}}, t) \tag{1.2.4}$$

Following the same steps with the second term on the right side of Eq. (1.2.2), we obtain

$$\sum_{N_{\text{U}}=0}^{\infty} \sum_{N_{\text{M}}=0}^{\infty} v\, N_{\text{U}}(N_{\text{M}}+1) P(N_{\text{U}}, N_{\text{M}}+1, t) = \sum_{N_{\text{U}}=0}^{\infty} \sum_{N_{\text{M}}=0}^{\infty} v\, N_{\text{U}} N_{\text{M}} P(N_{\text{U}}, N_{\text{M}}, t) \tag{1.2.5}$$

which, replacing Eqs. (1.2.4) and (1.2.5) in Eq. (1.2.2) and simplifying, leads to

$$\frac{\partial}{\partial t} \sum_{N_{\text{U}}=0}^{\infty} \sum_{N_{\text{M}}=0}^{\infty} N_{\text{U}} P(N_{\text{U}}, N_{\text{M}}, t) = - \sum_{N_{\text{U}}=0}^{\infty} \sum_{N_{\text{M}}=0}^{\infty} \Lambda(t)\, N_{\text{U}} P(N_{\text{U}}, N_{\text{M}}, t) \tag{1.2.6}$$

Finally, using the definition of the mean number of undecided individuals in Eq. (1.2.1), and the fact that $\Lambda(t)$ is a linear combination of Heaviside functions and thus a piecewise function taking constant values, we obtain

$$\dot{U}(t) = -\Lambda(t)\, U(t). \tag{1.2.7}$$

Following similar steps for the mean number of migrating individuals we get

$$\dot{M}(t) = \Lambda(t)\, U(t) - v\, M(t) \tag{1.2.8}$$

and, because $\dot{U}(t) + \dot{M}(t) + \dot{X}(t) = 0$,

$$\dot{X}(t) = v\, M(t). \tag{1.2.9}$$

We test the validity of this deterministic approximation across a range of parameter values and initial population sizes, obtaining excellent agreement between the mean population sizes obtained from an ensemble of realizations of the stochastic, individual-based simulations and the integration of the system of deterministic equations (1.2.7)-(1.2.9) (Fig. S2).

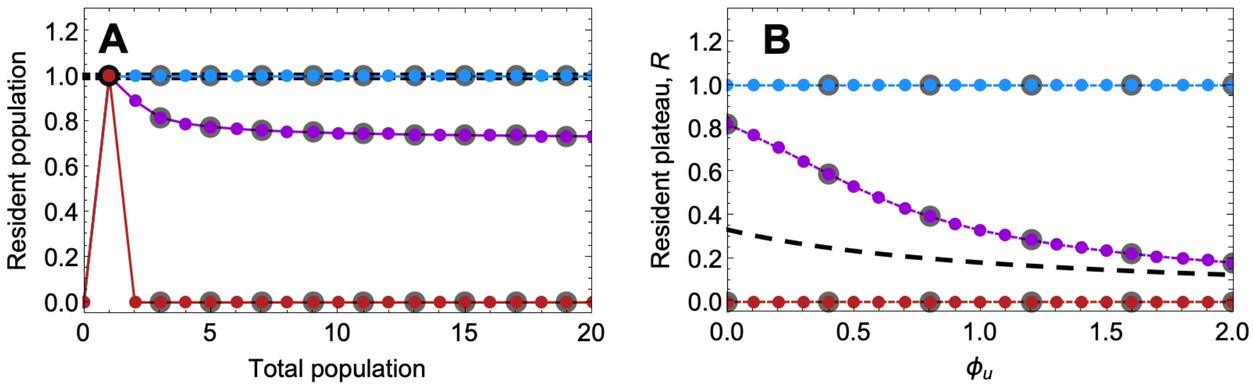

**Fig. S2:** Validation of the deterministic approximation using simulations of the individual-based stochastic dynamics. A) Scaled resident subpopulation versus the scaled total population (same as Fig. 3B in the main text). B) Scaled resident plateau versus $\phi_U$ (same as Fig. 4B in the main text). In both panels, the larger gray symbols show the average number of residents obtained from $10^4$ realizations of the stochastic dynamics. Parameter values (unless varied in the horizontal axes): $\theta = 0.4$; $\phi_M = 0.8$; $\phi_U = 0.1$; $U_0 = 6$; $(\lambda_E, \lambda_S) = (0.0, 0.2)$ red; $(0.4, 0.2)$ light purple; $(0.4, 0.0)$ blue; $(15, 15)$ dark purple. We use scaled parameters (see Section 1.3) to facilitate the comparison with the deterministic approximation.

#### 1.3 Analytical calculations

To simplify the analysis of the set of coupled ordinary differential equations in Eqs. (1.2.7)-(1.2.9), we define a series of dimensionless, scaled parameters and variables. We use these quantities only to obtain the patterns of partial migration for different model parameterizations, which we will characterize by the number of residents $R_{S,E}$. In our calculations, we will use different subscripts to indicate scenarios in which migration is driven only by one type of cues or by both: $R_{S,E}$ is the resident subpopulation when both types of cues matter, while $R_E$ and $R_S$ refer to the resident subpopulation when migration is driven only by environmental or social cues, respectively. Once we obtain the resident counts, we will transform these quantities back into their dimensional form. In the main text, we present the results using the original dimensional parameters and variables to make it explicit how different cues shape the resident subpopulation in our model.

We scale all mean population sizes by the bad-season carrying capacity, $\Theta_E/\psi$, to define dimensionless mean population sizes

$$u = \psi \frac{U}{\Theta_E} \qquad m = \psi \frac{M}{\Theta_E} \qquad x = \psi \frac{X}{\Theta_E} \qquad u_0 = \psi \frac{U_0}{\Theta_E} \qquad r = \psi \frac{R}{\Theta_E}. \tag{1.3.1}$$

Second, we scale time by the $M$-to-$X$ transition rate $v$,

$$\tilde{t} = v\, t \qquad \lambda = \frac{\Lambda}{v} \qquad \lambda_E = \frac{\Lambda_E}{v} \qquad \lambda_S = \frac{\Lambda_S}{v}. \tag{1.3.2}$$

Finally, we define the threshold ratio

$$\theta = \psi \frac{\Theta_S}{\Theta_E}. \tag{1.3.3}$$

These scalings lead to the simplified model equations we analyzed in the main text

$$\dot{u}(t) = -\lambda(t)u(t), \tag{1.3.4}$$
$$\dot{m}(t) = \lambda(t)u(t) - m(t), \tag{1.3.5}$$
$$\dot{x}(t) = m(t). \tag{1.3.6}$$

The scaled $U$-to-$M$ transition rate in Eqs. (1.3.4) and (1.3.5) is given by

$$\lambda(t) = \lambda_{\mathrm{S}} H(c_{\mathrm{S}}(t) - \theta) + \lambda_{\mathrm{E}} H(c_{\mathrm{E}}(t) - 1) \tag{1.3.7}$$

where $c_{\mathrm{S}}$ and $c_{\mathrm{E}}$ are the social and environmental cues after scaling the mean sub-population sizes,

$$c_{\mathrm{S}}(t) = \phi_{\mathrm{U}} u(t) + \phi_{\mathrm{M}} m(t) \tag{1.3.8}$$
$$c_{\mathrm{E}}(t) = u(t). \tag{1.3.9}$$

Next, using these scaled equations, we obtain model predictions for the resident subpopulation. In our model, the resident subpopulation comprises undecided individuals that do not transition to the migrating state when the intensity of cues is high (and hence $\lambda(t) \neq 0$). The scaled number of residents is given by

$$r_{\mathrm{S,E}} = u(\tau), \tag{1.3.10}$$

where $\tau$ is the time such that $\lambda(\tau) = 0$.

Because $\lambda$ is the linear combination of two theta-Heaviside functions

$$\tau = \max\left(\tau_{\mathrm{E}},\, \tau_{\mathrm{S}}\right), \tag{1.3.11}$$

where $\tau_{\mathrm{E}}$ and $\tau_{\mathrm{S}}$ are the times at which each of the theta-Heaviside functions in Eq. (1.3.7) becomes equal to zero. The values of these two times can be obtained from the implicit equations

$$u(\tau_{\mathrm{E}}) = 1 \tag{1.3.12}$$
$$\phi_{\mathrm{U}}\, u(\tau_{\mathrm{S}}) + \phi_{\mathrm{M}}\, m(\tau_{\mathrm{S}}) = \theta \tag{1.3.13}$$

and depending on the model parameterization, $\tau_{\mathrm{E}} < \tau_{\mathrm{S}}$ or $\tau_{\mathrm{E}} > \tau_{\mathrm{S}}$.

##### 1.3.1 Special case: Non-spatial limit.

In the non-spatial limit, individuals do not spend time in the migrating state and transition directly from the undecided to the departed state. Therefore, $m(t) = 0$ for every time $t$, which is equivalent to considering $\phi_{\mathrm{M}} = 0$. In this limit, we can obtain $\tau$ from Eqs. (1.3.12) and (1.3.13) because both social and environmental cues only depend on the subpopulation of undecided individuals,

$$\tau = \begin{cases} \tau_{\mathrm{E}} & \text{if } \theta > \phi_{\mathrm{U}} \\ \tau_{\mathrm{S}} & \text{otherwise} \end{cases} \tag{1.3.14}$$

The resident subpopulation will thus be equal to the threshold that determines the value of $\tau$. That is,

$$r_{\mathrm{S,E}} = \begin{cases} 1 & \text{if } \tau = \tau_{\mathrm{E}} \\ \dfrac{\theta}{\phi_{\mathrm{U}}} & \text{if } \tau = \tau_{\mathrm{S}} \end{cases} \tag{1.3.15}$$

equivalently,

$$r_{\mathrm{S,E}} = \min\left(1, \frac{\theta}{\phi_{\mathrm{U}}}\right), \tag{1.3.16}$$

or in terms of the original variables and parameters

$$R_{\mathrm{S,E}} = \min\left(\frac{\Theta_{\mathrm{E}}}{\psi}, \frac{\Theta_{\mathrm{S}}}{\phi_{\mathrm{U}}}\right). \tag{1.3.17}$$

##### 1.3.2 Special case: Only social cues determine migration calculation of $r_{\mathrm{S}}$

For completeness, and because this result will later be used to derive bounds for the general case where migration is driven by both social and environmental cues, we first consider the biologically implausible scenario in which migration is triggered solely by social cues.

In this special case, $\tau = \tau_{\mathrm{S}}$ and the model has three dynamical regimes. First, for $t \in [0, \tau]$ the sub-population sizes change according to

$$\dot{u}(t) = -\lambda_{\mathrm{S}}\, u(t) \tag{1.3.18}$$

$$\dot{m}(t) = \lambda_{\mathrm{S}}\, u(t) - m(t) \tag{1.3.19}$$

$$\dot{x}(t) = m(t) \tag{1.3.20}$$

Second, once the social cues reach the threshold $\theta$ at time $t = \tau$, the $U$-to-$M$ transition stops, which establishes the sub-population of residents. Migrants, however, continue to transition to the departed state. Finally, once the population of migrants reaches zero, the dynamics stops.

Because we are interested in determining the resident sub-population size, we focus on the end of the first dynamical regime $t = \tau$. Integrating Eqs. (1.3.18)-(1.3.20), evaluating the solutions at $t = \tau$ and replacing the expressions in the condition for the $U$-to-$M$ transition to stop given by Eq. (1.3.13) we obtain an implicit expression for $\tau$

$$\phi_{\mathrm{U}}\, u_0\, \mathrm{e}^{-\lambda_{\mathrm{S}}\tau} + \phi_{\mathrm{M}}\, u_0 \frac{\lambda_{\mathrm{S}}}{1-\lambda_{\mathrm{S}}}\left(\mathrm{e}^{-\lambda_{\mathrm{S}}\tau} - \mathrm{e}^{-\tau}\right) = \theta \tag{1.3.21}$$

By definition, the scaled resident subpopulation size is given by (1.3.10), which, in this special case, can be further resolved via integration to yield:

$$r_{\mathrm{S}} = u_0\, \mathrm{e}^{-\lambda_{\mathrm{S}}\tau(u_0)}, \tag{1.3.22}$$

where we have explicitly written the dependence of $\tau$ on the initial number of undecided individuals. Because we are interested in the resident subpopulation in the limit where it is independent of the initial population size, we further consider the limit $u_0 \to \infty$ in Eq. (1.3.22),

$$r_{\rm S} = \lim_{u_0 \to \infty} u_0 \, {\rm e}^{-\lambda_{\rm S} \tau(u_0)}. \tag{1.3.23}$$

The limit in Eq. (1.3.23) exists and converges to a finite value (Rossine et al., 2020), so we can compute it by applying L'Hôpital,

$$r_{\rm S} = \lim_{u_0 \to \infty} \frac{{\rm e}^{-\lambda_{\rm S} \tau(u_0)}}{\lambda_{\rm S} \, \tau'(u_0)}, \tag{1.3.24}$$

To calculate $\tau'(u_0)$, we differentiate both sides of Eq. (1.3.21) with respect to $u_0$, applying the chain rule when needed. We obtain,

$$A \, {\rm e}^{-\lambda_{\rm S} \tau(u_0)} - \phi_{\rm M} \, \lambda_{\rm S} \, {\rm e}^{-\tau(u_0)} + u_0 \, \lambda_{\rm S} \left[ \phi_{\rm M} \, {\rm e}^{-\tau(u_0)} - A \, {\rm e}^{-\lambda_{\rm S} \tau(u_0)} \right] \tau'(u_0) = 0 \tag{1.3.25}$$

where $A \equiv \left[ \phi_{\rm U}(1 - \lambda_{\rm S}) + \phi_{\rm M} \lambda_{\rm S} \right]$. Using Eq. (1.3.21) and arraging terms in Eq. (1.3.25) we obtain

$$\tau'(u_0) = \frac{\theta(1 - \lambda_{\rm S})}{u_0^2 \lambda_{\rm S} \left( A {\rm e}^{-\lambda_{\rm S} \tau(u_0)} - \phi_{\rm M} {\rm e}^{-\tau(u_0)} \right)}. \tag{1.3.26}$$

Next, we insert Eq. (1.3.26) in Eq. (1.3.24) and get

$$r_{\rm S} = \frac{1}{\theta(1 - \lambda_{\rm S})} \left[ A \, r_{\rm S}^2 - \phi_{\rm M} r_{\rm S} \, q \right] \tag{1.3.27}$$

where we have already computed the limits, used the definition of $r_{\rm S}$ from Eq. (1.3.23), and defined $q \equiv \lim_{u_0 \to \infty} u_0 {\rm e}^{-\tau(u_0)}$.

Eq. (1.3.27) is a quadratic equation in $r_{\rm S}$ with a second unknown $q$. One of the roots is $r_{\rm S} = 0$. To obtain the second root, we need another equation to solve together with Eq. (1.3.27) and calculate $r_{\rm S}$ and $q$. We obtain this second equation taking the limit $u_0 \to \infty$ in Eq. (1.3.21),

$$q = \frac{A \, r_{\rm S} - \theta(1 - \lambda_{\rm S})}{\phi_{\rm M} \, \lambda_{\rm S}}. \tag{1.3.28}$$

Finally, replacing $q$ from Eq. (1.3.28) into (1.3.27), we obtain the resident sub-population

$$r_{\rm S} = \frac{\theta(1 - \lambda_{\rm S})}{\phi_{\rm U}(1 - \lambda_{\rm S}) + \phi_{\rm M} \lambda_{\rm S}}, \tag{1.3.29}$$

which in terms of dimensional quantities is

$$R_{\rm S} = \frac{\Theta_{\rm S}(v - \Lambda_{\rm S})}{\phi_{\rm U}(v - \Lambda_{\rm S}) + \phi_{\rm M} \Lambda_{\rm S}}, \tag{1.3.30}$$

##### 1.3.3 General case: Both social and environmental cues influence migration; calculation of upper and lower bounds for $r_{\mathbf{S,E}}$

In this section, we derive some conditions for the existence of a resident subpopulation and obtain a lower bound for this subpopulation size in the limit of infinitely large total population size $u_0 \to \infty$.

Depending on whether the environmental or the social cues hit their respective threshold first, we can distinguish two cases in this analysis.

**Case A.-** $\tau_{\rm E} > \tau_{\rm S}$.

The end of the $U$-to-$M$ transition is determined by the environmental cues and $\tau = \tau_{\rm E}$. Because the environmental cues only depend on the undecided subpopulation, this case is equivalent to the non-spatial limit considered in Section 1.3.1 and $r_{\rm S,E} = 1$.

**Case B.-** $\tau_{\rm E} < \tau_{\rm S}$.

In this case, $u(\tau_{\rm S}) < 1$ because otherwise environmental cues would still stress undecided individuals and $\tau_{\rm S}$ would not be larger than $\tau_{\rm E}$. To obtain the expected number of residents for this case, we divide the migration into two phases. During the first phase, $t \in [0, \tau_{\rm E}]$, both environmental and social cues drive the $U$-to-$M$ transition and $\lambda = \lambda_{\rm S} + \lambda_{\rm E}$. During the second phase, $t \in [\tau_{\rm E}, \tau_{\rm S}]$, the environmental cues do not play any role in the dynamics and $\lambda = \lambda_{\rm S}$.

- *First phase:* $t \in [0, \tau_E]$. We integrate Eqs. (1.3.4) and (1.3.5) with initial condition $u(0) = u_0 > 1$, $m(0) = 0$. Notice that $u_0 > 1$ is a necessary condition to initiate the migration. We further set $\lambda = \lambda_{\rm S} + \lambda_{\rm E}$ because both the social and the environmental cues are above their thresholds. The dimensionless population size for each state at time $\tau_{\rm E}$ is

$$u(\tau_{\rm E}) = u_0\, {\rm e}^{-(\lambda_{\rm S}+\lambda_{\rm E})\tau_{\rm E}} \tag{1.3.31}$$
$$m(\tau_{\rm E}) = u_0 \frac{\lambda_{\rm S}+\lambda_{\rm E}}{1-(\lambda_{\rm S}+\lambda_{\rm E})}\left({\rm e}^{-(\lambda_{\rm S}+\lambda_{\rm E})\tau_{\rm E}} - {\rm e}^{-\tau_{\rm E}}\right). \tag{1.3.32}$$

  Solving Eq. (1.3.31) for $u(\tau_{\rm E}) = 1$, we get

$$\tau_{\rm E} = \frac{\ln u_0}{\lambda_{\rm E}+\lambda_{\rm S}}, \tag{1.3.33}$$

  which we can insert into Eq. (1.3.32) to calculate the size of the migrating subpopulation at time $\tau_{\rm E}$

$$m(\tau_{\rm E};\, u_0) = \frac{\lambda_{\rm S}+\lambda_{\rm E}}{1-(\lambda_{\rm S}+\lambda_{\rm E})}\left(1-(\psi u_0)^{1-\frac{1}{\lambda_{\rm E}+\lambda_{\rm S}}}\right). \tag{1.3.34}$$

  In the limit $u_0 \to \infty$, Eq. (1.3.34) yields

$$\lim_{u_0\to\infty} m(\tau_{\rm E};\, u_0) = \begin{cases} \infty & \text{if} \quad \lambda_{\rm S}+\lambda_{\rm E} > 1 \\ \frac{\lambda_{\rm S}+\lambda_{\rm E}}{1-(\lambda_{\rm S}+\lambda_{\rm E})} & \text{if} \quad \lambda_{\rm S}+\lambda_{\rm E} < 1 \end{cases} \tag{1.3.35}$$

- *Second phase:* $t \in [\tau_E, \tau_S]$. If $\lambda_{\rm S} + \lambda_{\rm E} > 1$, the migrating subpopulation at time $\tau_{\rm E}$ is infinite, and it will maintain the intensity of the social cues above its threshold indefinitely. As a result, all the remaining undecided individuals, $u(\tau_{\rm E}) = 1$, will transition to the migrating state during this second phase and, consequently, $r_{\rm S,E} = 0$. Therefore we continue the analysis under the assumption that $\lambda_{\rm S} + \lambda_{\rm E} < 1$.

  To simplify the notation, we shift the origin of time and set $\tau_{\rm E} = 0$. With this choice, integrating the second phase of the migration reduces to integrating Eqs. (1.3.4) and (1.3.5) with initial conditions $u(0) = 1$ and $m(0) = m(\tau_{\rm E})$ given by Eq. (1.3.35) in the limit $u_0 \to \infty$. We get

$$u(t) \;=\; {\rm e}^{-\lambda_{\rm S} t} \tag{1.3.36}$$

$$m(t) \;=\; \left(\frac{\lambda_{\rm S}+\lambda_{\rm E}}{1-(\lambda_{\rm S}+\lambda_{\rm E})} - \frac{\lambda_{\rm S}}{1-\lambda_{\rm S}}\right){\rm e}^{-t} + \frac{\lambda_{\rm S}}{1-\lambda_{\rm S}}{\rm e}^{-\lambda_{\rm S} t} \tag{1.3.37}$$

To get the size of the resident subpopulation, $r_{\rm S,E} = u(\tau_{\rm s})$, we would first have to calculate $\tau_{\rm S}$ from Eq.(1.3.13), which is not possible. However, we can calculate a lower bound for $r_{\rm S,E}$ and subsequently we will also calculate an upper bound.

**Lower bound.** To obtain this lower bound, we first use that $\exp(-t) < \exp(-\lambda_{\rm S} t)\ \forall t > 0$, because $\lambda_{\rm S} < \lambda_{\rm S} + \lambda_{\rm E} < 1$. Therefore, replacing $\exp(-t)$ by $\exp(-\lambda_{\rm S} t)$ in Eq. (1.3.37), we get

$$m(t) < \frac{\lambda_{\rm S}+\lambda_{\rm E}}{1-(\lambda_{\rm S}+\lambda_{\rm E})}{\rm e}^{-\lambda_{\rm S} t} \tag{1.3.38}$$

which we can use in Eq. (1.3.13) to get

$$\phi_{\rm U}\, r_{\rm S,E} + \phi_{\rm M}\, \frac{\lambda_{\rm S}+\lambda_{\rm E}}{1-(\lambda_{\rm S}+\lambda_{\rm E})} r_{\rm S,E} > \theta, \tag{1.3.39}$$

where we used the fact that $\exp(-\lambda_{\rm S}\tau_{\rm S}) = u(\tau_{\rm S}) \equiv r_{\rm S,E}$, Finally, solving for $r_{\rm S,E}$, we find the lower bound for the resident subpopulation to be given by

$$r_{\rm S,E} \;>\; \theta \frac{1-(\lambda_{\rm S}+\lambda_{\rm E})}{\phi_{\rm U}[1-(\lambda_{\rm S}+\lambda_{\rm E})] + \phi_{\rm M}(\lambda_{\rm S}+\lambda_{\rm E})} \tag{1.3.40}$$

which in terms of dimensional quantities is

$$R_{\rm S,E} \;>\; \Theta_{\rm S} \frac{v-(\Lambda_{\rm S}+\Lambda_{\rm E})}{\phi_{\rm U}[v-(\Lambda_{\rm S}+\Lambda_{\rm E})] + \phi_{\rm M}(\Lambda_{\rm S}+\Lambda_{\rm E})}. \tag{1.3.41}$$

.

**Upper bound.** Next, we will try to also obtain an upper bound for $r_{\rm S,E}$. Specifically, we will show that $r_{\rm S}$ is an upper bound for $r_{\rm S,E}$ when $r_{\rm S} < 1$. We work under the assumption that $r_{\rm S} < 1$ because when $r_{\rm S} > 1$ the upper-bound for the resident subpopulation when both environmental and social cues influence the decision-making is 1. This is so because environmental cues will drive migration for as long as $u(t) > 1$ and thus make $r_{\rm S,E} \leq 1$. In this section, we prove that $r_{\rm S}$ is a stricter upper bound for $r_{\rm S,E}$ when $r_{\rm S} < 1$.

To prove this result, we depart from the condition for the migration to stop in Eq. (1.3.13)

$$\phi_{\rm U}\, u(\tau_{\rm S}) + \phi_{\rm M}\, m(\tau_{\rm S}) = \theta. \tag{1.3.42}$$

Replacing $u(\tau_S)$ and $m(\tau_S)$ with their form from Eqs. (1.3.36) and (1.3.37), we obtain

$$e^{-\lambda_S \tau_S} + \frac{\phi_M}{\phi_U}\left(Ae^{-\tau_S} + \frac{\lambda_S}{1-\lambda_S}e^{-\lambda_S \tau_S}\right) = \frac{\theta}{\phi_u}, \tag{1.3.43}$$

where $A > 0$ depends on $\lambda_S$ and $\lambda_E$. Using the definition of $r_{S,E}$, we get

$$r_{S,E} + \frac{\phi_M}{\phi_U}\left(Ae^{-\tau_S} + \frac{\lambda_S}{1-\lambda_S}r_{S,E}\right) = \frac{\theta}{\phi_u}, \tag{1.3.44}$$

which, defining $k \equiv \theta/\phi_u$ and $\phi \equiv \phi_m/\phi_u$ gives

$$r_{S,E} = \frac{\left[k - \phi\, Ae^{-\tau_S}\right](1-\lambda_S)}{1-\lambda_S+\phi\lambda_S} \tag{1.3.45}$$

Using Eq. (1.3.29) for the resident subpopulation size when migration is driven solely by social cues, we can further write

$$r_{S,E} = \frac{k - \phi\, Ae^{-\tau_S}}{k} r_S < r_S \tag{1.3.46}$$

because $\phi_M\, Ae^{-\tau_S}/\theta > 0$.

### 2 Dimorphic population

#### 2.1 Model extension

We extend the model to describe the onset of migration in a dimorphic population with small and large individuals. We make the simplifying assumption that body size does not determine the values of $v$, $\Lambda_E$, and $\Lambda_S$. Instead, we introduce heterogeneity through differences in sensitivity thresholds, in the parameters that weigh the contributions of migrating and undecided individuals to these cues, and in per-capita resource demands $\psi$. Moreover, because migration onset occurs on short time scales in which changes in individual body size are likely negligible, we also assume that small individuals do not become large. For notational clarity, we implement heterogeneity directly in the deterministic model, but it is straightforward to develop a stochastic model for a dimorphic population and obtain its deterministic limit following the steps in Section 1.2.

First, we split the dynamics of undecided and migrating individuals into two different equations for each class, one for each possible body size. We do not split the dynamics of the departed individuals into body-size classes because it is a redundant equation that we will not use in our analysis. This division of the undecided and migrating individuals in terms of their body size leads to a model consisting of five coupled ordinary differential equations

$$\dot{U}_\alpha(t) = -\Lambda_\alpha(t)\, U_\alpha(t) \tag{2.1.1}$$

$$\dot{M}_\alpha(t) = \Lambda_\alpha(t)\, U_\alpha(t) - vM_\alpha(t) \tag{2.1.2}$$

$$\dot{X}(t) = vM(t) \tag{2.1.3}$$

where the subscript $\alpha$ labels the body-size class. $\alpha = \{\ell, s\}$ for large and small individuals respectively. As we did for the monomorphic case, we consider an initial population with all individuals in the undecided state. To account for the body-size composition in the initial population, we introduce a parameter $f$ that gives the fraction of individuals belonging to the small body-size class. Mathematically, we can write this initial condition as $U_s(0) = f\,U_0$, $U_\ell(0) = (1-f)U_0$; furthermore, $M_s(0) = M_\ell(0) = X(0) = 0$.

Second, considering two different body sizes impacts the $U$-to-$M$ transition rate $\Lambda_\alpha(t)$ in (2.1.1)-(2.1.3). This transition rate is given by

$$\Lambda_\alpha(t) = \Lambda_S\, H\big(C_{S,\alpha}(t)\big) + \Lambda_E\, H\big(C_{E,\alpha}(t)\big) \tag{2.1.4}$$

where $\alpha = \{\ell,\, s\}$. Finally, we also allow individuals of different body sizes to potentially contribute to the concentration of cues differently

$$C_{E,\alpha}(t) \;=\; \sum_{\beta=\{\ell,s\}} \psi_\beta\, U_\beta(t) - \Theta_{E\alpha} \tag{2.1.5}$$

$$C_{S,\alpha}(t) \;=\; \sum_{\beta=\{\ell,s\}} \big(\phi_{u\beta} U_\beta(t) + \phi_{m\beta} M_\beta(t)\big) - \Theta_{S\alpha}. \tag{2.1.6}$$

As discussed in the main text, we assume in Eqs. (2.1.4)-(2.1.6) that smaller individuals have a higher environmental threshold than large individuals, i.e. $\Theta_{Es} > \Theta_{E\ell}$, because the former have lower resource demands and thus can tolerate higher levels of intraspecific competition. As a sensitivity analysis we repeat this procedure with $\Theta_{Es} = \Theta_{E\ell}$ (see Fig. S3). Using the same argument, larger individuals have higher resource demands and they contribute more strongly to intraspecific competition and environmental cues. This difference in resource demands means that $\psi_\ell > \psi_s$. As expected, $\Theta_{Es} > \Theta_{E\ell}$ together with $\psi_\ell > \psi_s$ make the bad-season carrying capacity correlate negatively with individual body size.

#### 2.2 Definition of probability of migrating, $\mathcal{P}$, and skew in body size among resident subpopulation, $\mathcal{B}$

We define two metrics to quantify the impact of population heterogeneity on patterns of partial migration. First, we measure the skew in the residents relative to the initial composition of the population as the difference between the frequency of one body size in the resident subpopulation and its frequency in the initial population:

$$\mathcal{B}_\alpha = \frac{R_\alpha}{R} - \frac{U_\alpha(0)}{U(0)} \tag{2.2.1}$$

where $\alpha = \{\ell,\, s\}$, and $R = R_s + R_\ell$ is the total number of residents. Notice that $\mathcal{B}_s = -\mathcal{B}_\ell$ by definition.

Second, we measure the probability of migrating, an individual-level measure of partial migration that quantifies how likely each individual is to migrate, depending on its body size. This probability is given by

$$\mathcal{P}_\alpha = 1 - \frac{R_\alpha}{U_\alpha(0)} \tag{2.2.2}$$

with $\alpha = \{\ell, \mathrm{s}\}$.

#### 2.3 Analytical calculations for $\Lambda_{\mathrm{Es}} = \Lambda_{\mathrm{E}\ell}$

In this limit, the $U$-to-$M$ transition rates simplify to

$$\Lambda_\ell(t) = \Lambda_{\mathrm{E}}\, H\left(\sum_{\beta=\{\ell,\mathrm{s}\}} \psi_\beta\, U_\beta(t) - \Theta_{\mathrm{E}\ell}\right), \tag{2.3.1}$$

$$\Lambda_{\mathrm{s}}(t) = \Lambda_{\mathrm{E}}\, H\left(\sum_{\beta=\{\ell,\mathrm{s}\}} \psi_\beta\, U_\beta(t) - \Theta_{\mathrm{Es}}\right). \tag{2.3.2}$$

Considering these expressions for the $U$-to-$M$ transitions rates, because $\Theta_{\mathrm{Es}} > \Theta_{\mathrm{E}\ell}$ and $U_{\mathrm{s}}$ and $U_\ell$ are monotonically decreasing functions of time, $\Lambda_{\mathrm{s}}$ reaches zero before $\Lambda_\ell$. Moreover, because $\Lambda_{\mathrm{E}}$ is the same for both body-size classes, we can calculate the small-individual resident subpopulation size using the fact that $U_{\mathrm{s}}(t) = f\,U(t)$ and $U_\ell(t) = (1-f)U(t)$ and solving the argument of the Heaviside-theta function in Eq. (2.3.2). We get

$$R_{\mathrm{s}} = \frac{\Theta_{\mathrm{Es}}\, f}{f\psi_{\mathrm{s}} + (1-f)\psi_\ell} \tag{2.3.3}$$

To obtain the number of residents belonging to the large body-size class, we substitute the small body-size resident subpopulation from Eq. (2.3.3) in the argument of the Heaviside-theta function of Eq. (2.3.1) and get

$$R_\ell = \max\left(\frac{\Theta_{\mathrm{E}\ell} - \psi_{\mathrm{s}} R_{\mathrm{s}}}{\psi_\ell},\ 0\right). \tag{2.3.4}$$

Finally, we can obtain the total number of residents by combining these two results $R = R_{\mathrm{s}} + R_\ell$ (blue curve in Fig. 5A).

Substituting Eqs. (2.3.3) and (2.3.4) in the definition of $\mathcal{P}$ in Eq. (2.2.2), we obtain the probability of migrating for individuals belonging to each body-size class in a dimorphic population (light and dark blue curves in Fig. 5C, D). Because small individuals in our model have a higher environmental threshold than large individuals, $\Theta_{\mathrm{Es}} > \Theta_{\mathrm{E}\ell}$, and we are not considering the effect of social information on the decision of whether to migrate or not, we expect that small individuals should be less likely to migrate than large individuals in a dimorphic population. To prove this expectation, we calculate $\mathcal{P}_\ell - \mathcal{P}_{\mathrm{s}}$. This difference must be positive if our prediction is true. After some algebra, we obtain

$$\mathcal{P}_\ell - \mathcal{P}_{\mathrm{s}} = \frac{\Theta_{\mathrm{E}\ell} - \Theta_{\mathrm{Es}}}{(1-f)\psi_\ell} \tag{2.3.5}$$

which is positive because $\Theta_{\mathrm{E}\ell} > \Theta_{\mathrm{Es}}$.

### 3 Model supplementary figures

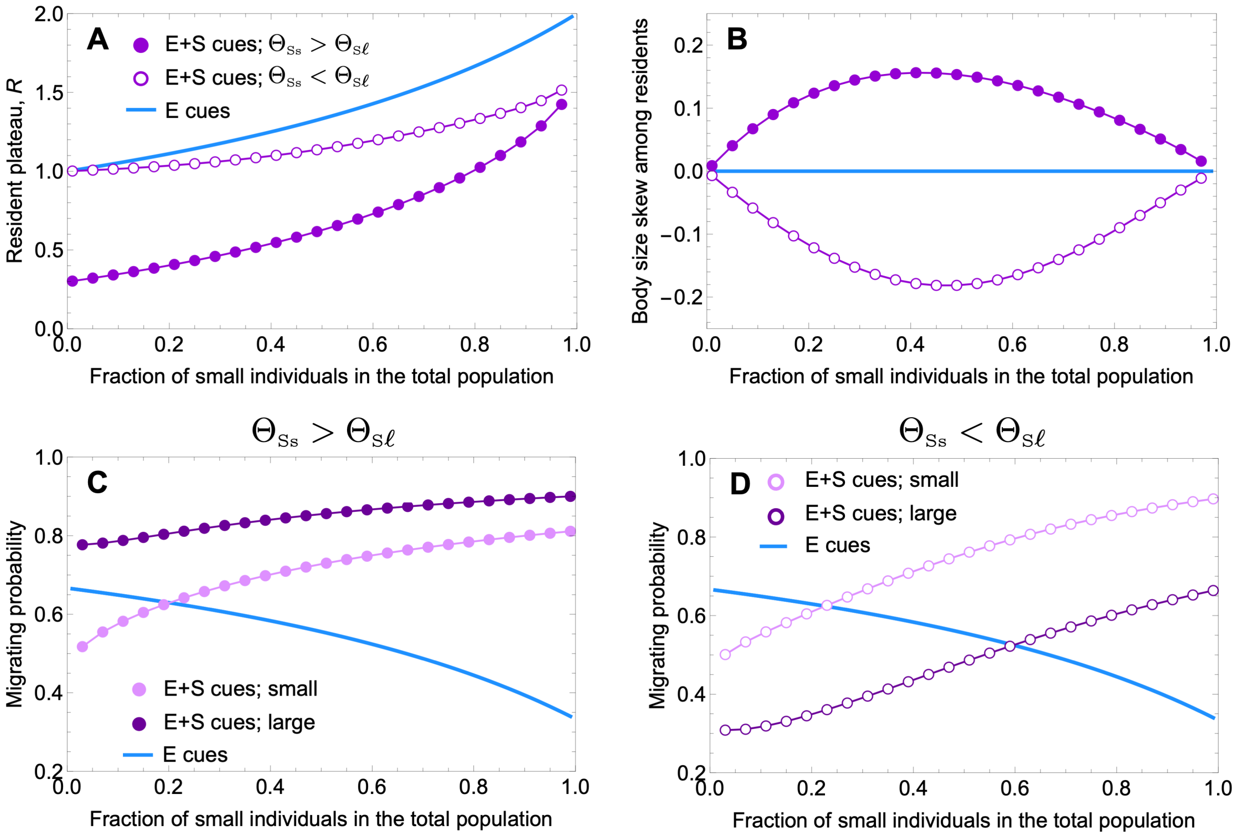

**Fig. S3:** Model predictions for heterogeneous populations with the same environmental threshold. (A) The resident plateau continues to depend on the proportion of small and large-bodied individuals in the population, as well as on the relationship between their social thresholds. (B) If social cues do not factor into the decision to migrate, the proportion of small individuals is the same between the resident subpopulation and the total population. When social cues contribute, small individuals are overrepresented in the resident subpopulation relative to the total population when $\Theta_{\mathrm{Ss}} > \Theta_{\mathrm{S}\ell}$ but are underrepresented in the resident subpopulation when $\Theta_{\mathrm{Ss}} < \Theta_{\mathrm{S}\ell}$. (C, D) These skews within the resident subpopulation translate into probabilities that individuals from each size class migrate: (C) small individuals are less likely to migrate than large individuals when $\Theta_{\mathrm{Ss}} > \Theta_{\mathrm{S}\ell}$, but (D) small individuals are more likely to migrate than large individuals when $\Theta_{\mathrm{Ss}} < \Theta_{\mathrm{S}\ell}$. Parameters: $U_0 = 3$; $\phi_{\mathrm{M}\ell} = 0.8$; $\phi_{\mathrm{Ms}} = 0.4$; $\phi_{\mathrm{U}\ell} = 0.1$; $\phi_{\mathrm{Us}} = 0.08$; $\Lambda_{\mathrm{S}} = 0.4$; $\Lambda_{\mathrm{E}} = 0.4$; $\psi_\ell = 1$; $\psi_{\mathrm{s}} = 0.5$; $v = 1$; $\Theta_{\mathrm{Es}} = \Theta_{\mathrm{E}\ell} = 1$; $\Theta_{\mathrm{Ss}} = 0.4$; $\Theta_{\mathrm{S}\ell} = 0.8$ or $\Theta_{\mathrm{S}\ell} = 0.2$. See Part I for definitions of bias metrics and probability of migrating.

### References

Gillespie, D. T. (1977). Exact stochastic simulation of coupled chemical reactions. *The journal of physical chemistry*, 81(25):2340–2361.

Rossine, F. W., Martinez-Garcia, R., Sgro, A. E., Gregor, T., and Tarnita, C. E. (2020). Eco-evolutionary significance of "loners". *PLoS Biology*, 18(3):e3000642.

Toral, R. and Colet, P. (2014). *Stochastic numerical methods: an introduction for students and scientists*. John Wiley & Sons.

Van Kampen, N. G. (1992). *Stochastic processes in physics and chemistry*, volume 1. Elsevier.
